# DNA binding reorganizes the intrinsically disordered C-terminal region of PSC in *Drosophila* PRC1

**DOI:** 10.1101/2020.06.02.130492

**Authors:** Jin Joo Kang, Denis Faubert, Jonathan Boulais, Nicole J. Francis

## Abstract

Polycomb Group (PcG) proteins regulate gene expression by modifying chromatin. A key PcG complex, Polycomb Repressive Complex 1 (PRC1), has two activities: a ubiquitin ligase activity for histone H2A, and a chromatin compacting activity. In *Drosophila*, the Posterior Sex Combs (PSC) subunit of PRC1 is central to both activities. The N-terminal homology region (HR) of PSC assembles into PRC1, including partnering with dRING to form the ubiquitin ligase for H2A. The intrinsically disordered C-terminal region of PSC (PSC-CTR) compacts chromatin, and inhibits chromatin remodeling and transcription *in vitro*. Both the PSC-HR and the PSC-CTR are essential *in vivo*. To understand how these two activities may be coordinated in PRC1, we used cross-linking mass spectrometry (XL-MS) to analyze the conformations of the PSC-CTR in PRC1 and how they change on binding DNA. XL-MS identifies interactions between the PSC-CTR and the core of PRC1, including between the PSC-CTR and PSC-HR. New contacts and overall more compacted PSC-CTR conformations are induced by DNA binding. Protein footprinting of accessible lysine residues in the PSC-CTR reveals an extended, bipartite candidate DNA/chromatin binding surface. Our data suggest a model in which DNA (or chromatin) follows a long path on the flexible PSC-CTR. Intramolecular interactions of the PSC-CTR detected by XL-MS can bring the high affinity DNA/chromatin binding region close to the core of PRC1 without disrupting the interface between the ubiquitin ligase and the nucleosome. Our approach may be applicable to understanding the global organization of other large IDRs that bind nucleic acids.

**Highlights:**

- An intrinsically disordered region (IDR) in Polycomb protein PSC compacts chromatin
- Cross-linking mass spectrometry (XL-MS) was used to analyze topology of the PSC IDR
- Protein footprinting suggests a bipartite DNA binding surface in the PSC IDR
- A model for the DNA-driven organization of the PSC IDR
- Combining XL-MS and protein footprinting is a strategy to understand nucleic acid binding IDRs

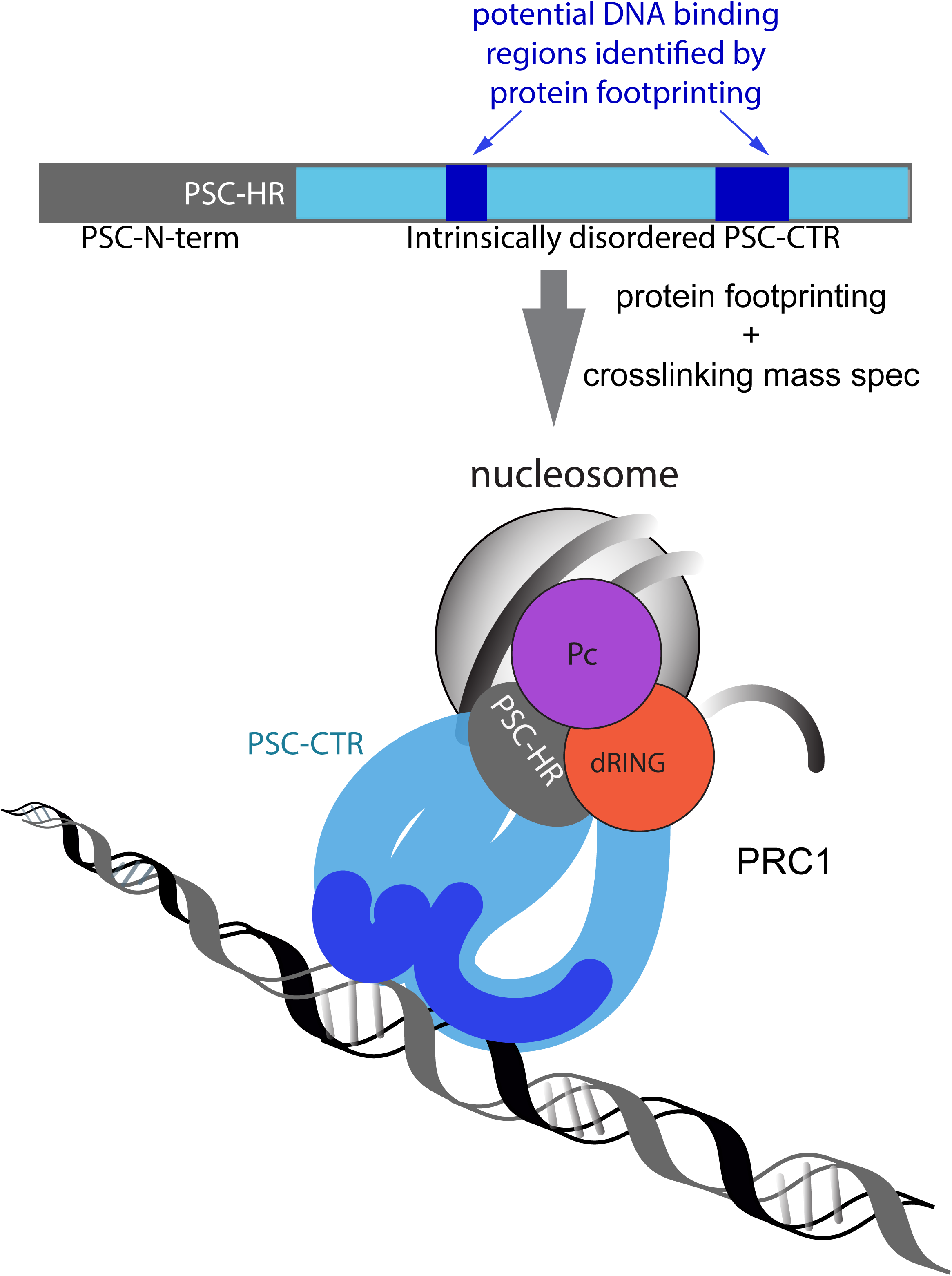

## Introduction

Polycomb Group (PcG)^1^ proteins are essential regulators of gene expression that assemble into multiprotein complexes. These complexes modify chromatin biochemistry and structure through histone post-translational modifications and non-covalent effects on chromatin organization[1-4]. Two main complexes, PRC1 and PRC2, are conserved through evolution and are central to PcG regulation of gene expression. PRC2 has a histone methyltransferase activity towards H3K27[5], and also interacts with RNA[6, 7]. PRC1 has an E3 ligase activity towards histone H2A, and also has powerful effects on chromatin organization[1-3].

In *Drosophila*, Posterior Sex Combs (PSC) is critical for both PRC1 activities. PSC contains conserved Ring and RAWUL (ubiquitin-like) domains that partner with dRING to ubiquitylate histones[8]. The N-terminal region of PSC harboring these domains has been termed the homology region (HR)[9]. PSC also has a ∼1150 amino acid (aa) C-terminal intrinsically disordered region (IDR) (hereafter referred to as the PSC-CTR) that binds tightly to chromatin and DNA[9-11]. The PSC-CTR mediates chromatin compaction[12], nucleosome bridging[13], and inhibition of transcription[9] and chromatin remodeling[9, 14], and is essential for PcG function *in vivo*[9]. In mammals, PRC1 is diversified into at least 6 subcomplexes[8]. Both the E3 ligase and chromatin organization functions of PRC1 are conserved, but they are partitioned among different subunits and paralogues[10, 15-17]. The conserved Sterile Alpha Motif (SAM) present in Polyhomeotic, another subunit of PRC1, also contributes to large-scale chromatin organization in *Drosophila* and mammals [18-20]. The effect of the Ph SAM is not considered in this work, which focuses on the PSC-CTR.

Nucleosome bridging, chromatin compaction, and persistent binding through DNA replication can be achieved by PRC1ΔPh, or by PSC or the PSC-CTR alone. For PSC alone, a ratio of 1 PSC per nucleosome is required[21], while for PRC1ΔPh, 1 complex can affect 3-4 nucleosomes[12]. We previously showed that the activity of PSC alone depends on both DNA binding and self-association of PSC, and suggested that self-association could also be important for the activities of PRC1ΔPh[13, 14]. However, PRC1ΔPh behaves as a monomer under conditions where PSC multimers are detected[14]. This raises the possibility that the function of the PSC-CTR may be different in the context of PRC1ΔPh versus when PSC (or the PSC-CTR) is analyzed alone.

We therefore sought to find additional methods to understand the organization and function of the PSC-CTR in the context of PRC1. We used lysine-based XL-MS to identify regions of PSC involved in protein-protein contacts, with an emphasis on intra-protein crosslinks that reflect (possible) folding of the PSC-CTR[22]. In addition, we used protein footprinting by chemical acetylation of accessible lysine residues to identify likely DNA/chromatin binding regions of the PSC-CTR. We combined the results from both methods to produce a global model for the topology of the PSC-CTR and how DNA binding changes its conformation in the context of PRC1ΔPh.

## Results & Discussion

### XL-MS analysis of PRC1ΔPh

To understand the basic topology of the PSC-CTR, how or whether it interacts with the core of PRC1, and how its conformation changes upon binding DNA, we subjected PRC1 to XL-MS. We used PRC1ΔPh, consisting of PSC, Polycomb (Pc), and dRING (Fig. 1), because this complex is biochemically tractable (sFig. 1A) and recapitulates the activities of PRC1 on chromatin and DNA[9, 12, 13, 23]. PRC1ΔPh is also an active E3 ubiquitin ligase for histone H2A (Seif et al.,in revision). PSC, particularly the CTR, is rich in lysines, which are distributed throughout the protein (Fig. 1C). We therefore used the lysine crosslinker BS^3^.

**Figure 1.**
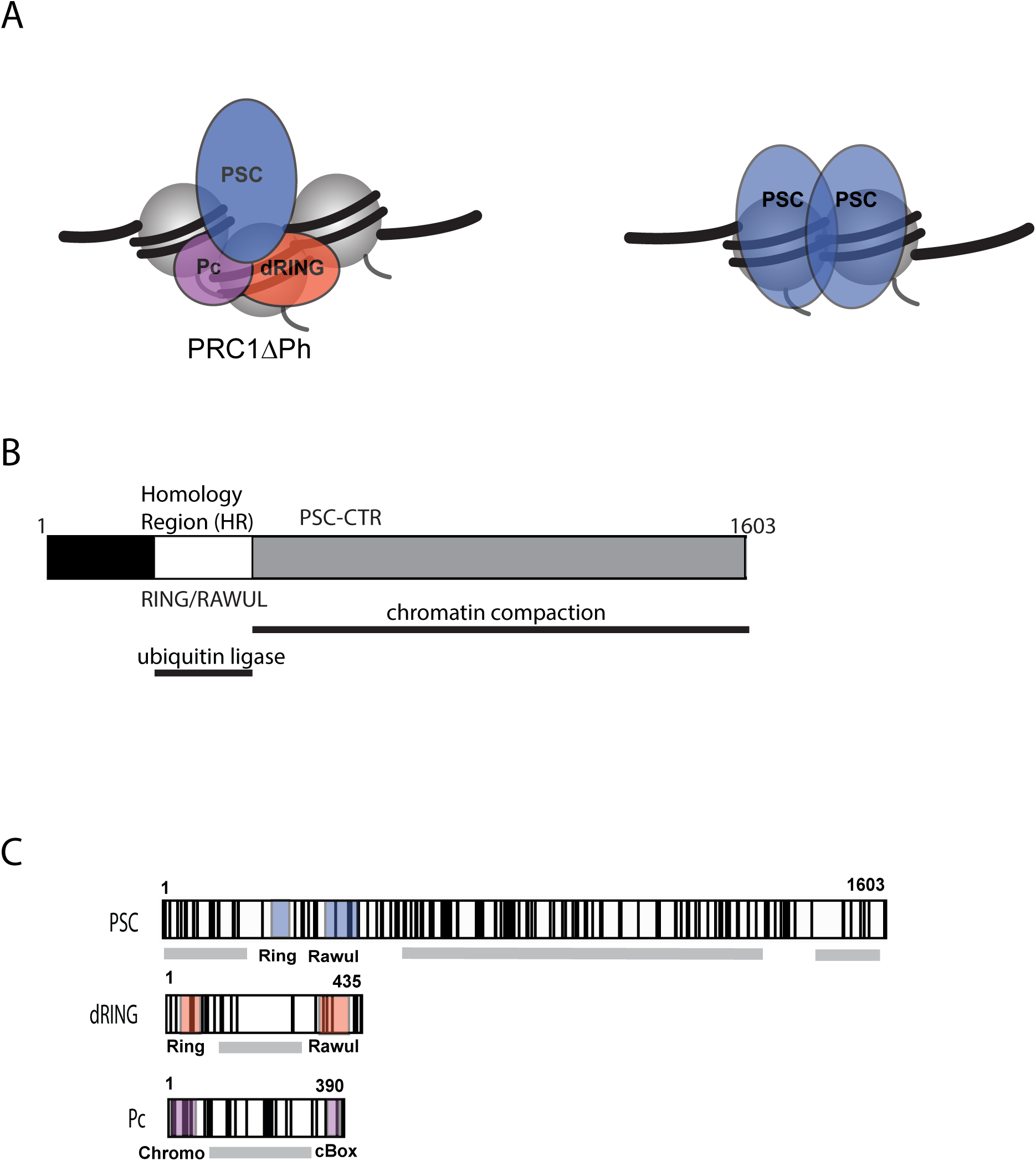
Assembly into PRC1 affects PSC activity. A. PRC1ΔPh (left) and PSC alone interact differently with chromatin. PRC1ΔPh has ubiquitin ligase activity and can compact chromatin at a ratio of 1 PRC1:3-4 nucleosomes. In contrast, PSC alone self-interacts and compacts chromatin at a ratio of 1:1 with nucleosomes. See Introduction. B. Schematic of PSC. Ubiquitin ligase activity and assembly into PRC1 requires the Homology Region (HR), while the disordered PSC-CTR binds DNA and compacts chromatin. C. Distribution of lysine residues in PRC1ΔPh. Black lines indicate the positions of lysines; gray bars indicate predicted disordered regions in each subunit, and known domains are shaded and labelled.

We carried out XL-MS on PRC1ΔPh alone, and PRC1ΔPh incubated with a 65 base pair DNA substrate. The PSC-CTR binds DNA without sequence specificity but with high affinity[10, 13], and high-affinity DNA binding is correlated with effects on chromatin[13]. We titrated BS^3^ and PRC1ΔPh concentrations and found that 20µg of PRC1ΔPh (76.6pmol), and 2mM BS^3^ consistently produced a major band at the size expected for a crosslinked monomeric complex (261kDa) (sFig. 1B). To enrich for complexes that are not hyper-crosslinked, we purified crosslinked PRC1ΔPh on glycerol gradients, selecting fractions with the visible high molecular weight band corresponding to PRC1ΔPh, and limited higher MW species and material trapped in the well (both of which are likely to be hyper-crosslinked) (sFig. 1C, D). We used MCX resin[24] to enrich crosslinked peptides based on charge, and analyzed the samples using an Orbitrap Fusion mass spectrometer.

We used three programs, pLink 2[25, 26], MeroX[27], and SIM-XL[28] to identify cross-linked peptides. Because we are most interested in the conformational dynamics of the PSC-CTR, we focused on intra-protein crosslinks of PSC, which were the majority of the crosslinks identified. To ensure that the crosslinking patterns observed are likely to be meaningful, we used stringent score cutoffs (20 for pLink, 120 for MeroX, and 4.0 for SIM-XL). We insisted that crosslinks be identified in at least one sample with a score above the threshold, and identified in all other samples (total of 4 samples without DNA and 3 with DNA), irrespective of score. The three programs identified largely non-overlapping crosslinks (Fig. 2A-E), although each program identified crosslinks with high quality spectra (Fig. 2F, sFig. 2). We therefore combined data from all three programs that passed our quality criteria. This resulted in a total of 75 unique crosslinks, 46 of which are intra-protein crosslinks in PSC. All of the identified crosslinks that passed our criteria are available in Supplementary Tables 1-6.

**Figure 2.**
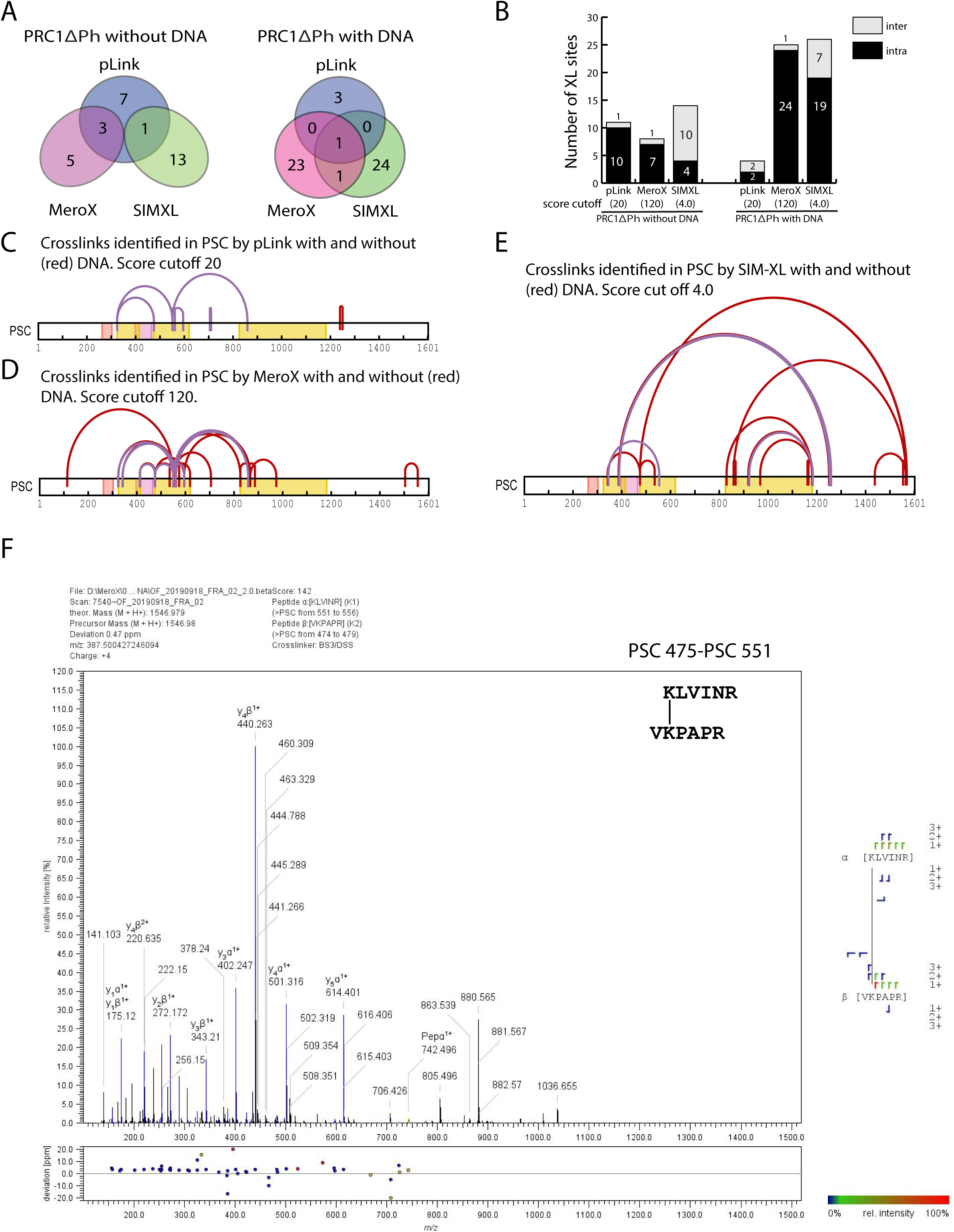
Identification of crosslinked peptides in PRC1ΔPh. A. Venn Diagram of overlap of crosslinking sites in PRC1ΔPh identified with high confidence by pLink, MeroX, and SIM-XL. B. Graph of inter- and intra-protein crosslinking sites of PRC1ΔPh identified in the absence or presence of DNA. Note that the scores are calculated differently by each algorithm so that different thresholds are used in each case, and they cannot be directly compared. C-E. Maps of intra-protein crosslinks identified in PSC by each program. F. Representative spectrum of crosslinked peptides identified by MeroX. See sFig. 2 for spectra generated by pLink and SIM-XL.

XL-MS data are usually validated by comparison with structural information. Although the majority of crosslinks obtained involve regions of PSC for which there is no structural information available, we obtained 8 inter- and intra-protein crosslinks within dRING that could be mapped onto existing structures of the Ring domains of human homologues of PSC and dRING. We built a homology model for *Drosophila* PSC/dRING, and mapped our crosslinking data onto it (sFig. 3). We measured the distance between Cα for each of the identified crosslinks. 4 out of 6 dRING intra-protein (sFig. 3A, sTable 6), and both dRING-PSC inter-protein (sFig. 3B, sTables 3, 4) crosslinks are below the suggested 30Å distance cutoff for valid lysine-lysine crosslinks[29]. The two remaining crosslinks were K22-S106 (31Å), and S77-S106 (30Å), which we discarded based on the shorter expected distance for serine than lysine. We conclude that our quality control criteria identify valid crosslinks.

### The PSC-CTR interacts with the N-terminal homology region

In the absence of DNA, we identified multiple intra-protein crosslinks within two stretches of sequence in PSC (Fig. 3A, sTable 1). We term these Protein Patch 1 (PP1) and Protein Patch 2 (PP2) (Fig. 2A). PP1 spans K324-K619, while PP2 spans K826-K1181. A small number of crosslinks between PP1 and PP2 are also observed, indicating that these regions may be close together in the absence of DNA.

**Figure 3.**
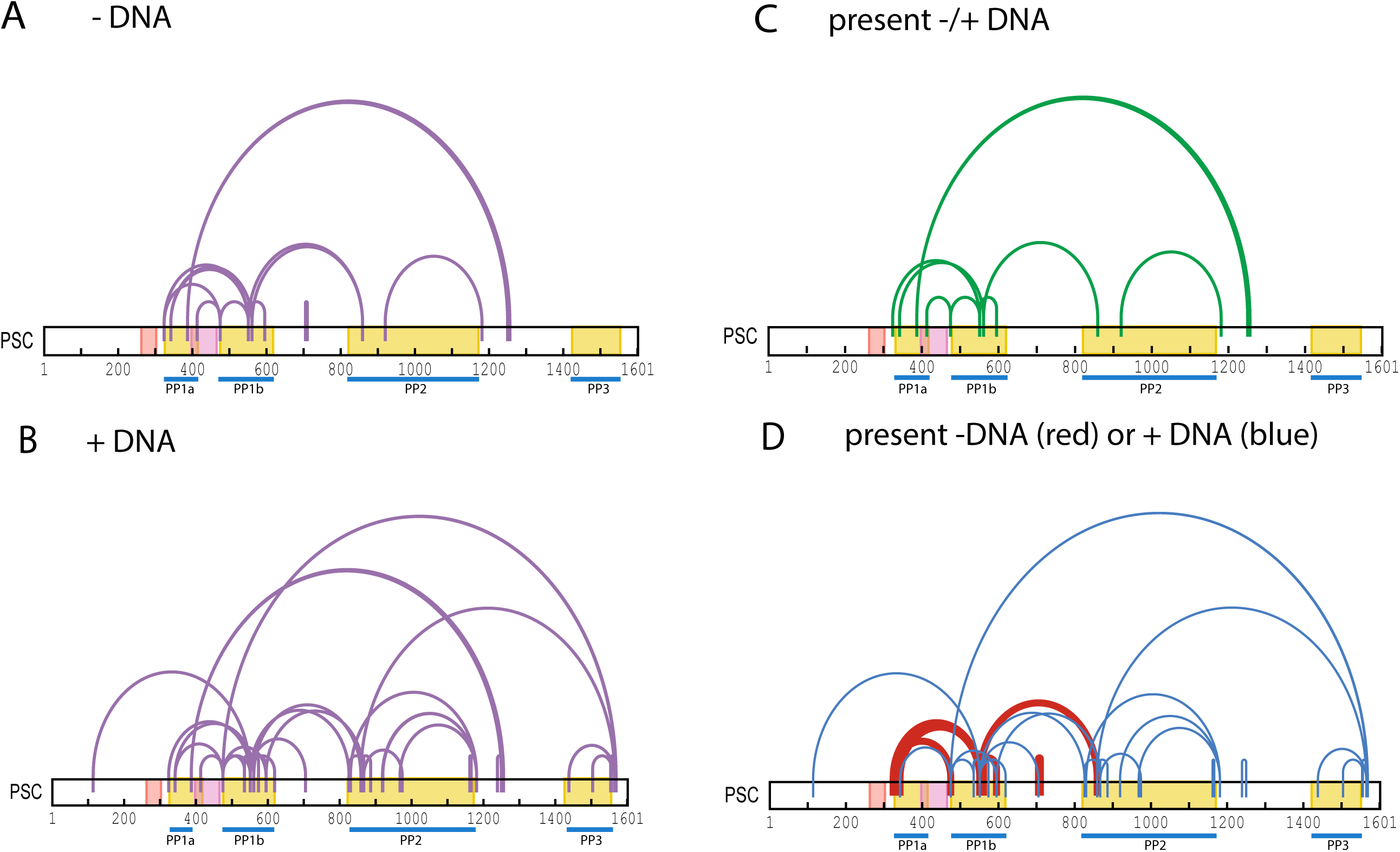
XL-MS analysis of PSC in the context of PRC1ΔPh with and without bound DNA. A, B. Intra-protein crosslinks identified in PSC in the absence (A) or presence (B) of DNA. C. Intra-protein crosslinks identified in *both* the absence and presence of DNA. D. Intra-protein crosslinks identified *only* in the absence (red) or presence (blue) of DNA.

PP1 can be split into PP1a (K324-K413), which is in the HR, and thus part of the core of PRC1, and PP1b (K475-K619), which is part of the PSC-CTR. PP1a and PP1b are largely separated by the RAWUL domain (K413 is within the RAWUL). Crosslinks between PP1a and PP1b thus reflect interactions between the PSC-CTR and the HR. Supplementary Fig. 3C shows the positions of K324 and K342, each of which forms crosslinks with PP1b, in the PSC-dRING Ring domain model. Notably, K324 is part of a previously identified conserved patch between *Drosophila* and mammalian PSC homologues (Y**K**LVPGL)[11] (sFig. 4). Finally, S387 within PP1a also crosslinks to K1250 and K1256, forming a second site of interaction between the PSC-CTR and the HR (Fig. 3A)

The crosslinking pattern within PP2 displays one interaction with PP1b as well as an intra-protein crosslink within PP2 itself (Fig. 3A).Although PP2 does not directly crosslink to the HR, the PP2-PP1b crosslink is expected to bring the CTR close to the HR. These results indicate the PSC-CTR folds onto itself, and onto the HR in the absence of DNA. Although the PSC-CTR is intrinsically disordered, the concentration of crosslinks in patches suggests a hierarchical organization in which each patch undergoes local interactions and interactions with other patches.

### The PSC-CTR compacts on binding DNA

In the presence of DNA, the number of intra-protein crosslinks is increased (18 without DNA, 35 with DNA) (Fig. 3B, sTable 1), while the majority of crosslinks observed in the absence of DNA are still present (Fig. 3C, sTable 2). Although the increase in crosslinks on DNA binding is the most prominent feature, there are also 9 crosslinks detected without DNA but not with DNA (Fig. 3D, sTable 1). In the presence of DNA, increased interactions are observed within PP1 and PP2 (Fig. 3B, D). The new crosslinks within PP1 and particularly PP2 could reflect more compact conformations of the PSC-CTR. Crosslinks between PP2 and PP1b are also increased, but there are still no crosslinks observed between PP2 and PP1a. The long-range crosslinks between K1250/K1256 and S387 (in PP1a) remain upon DNA binding.

In the presence of DNA, a new crosslinked patch, PP3, is also observed from K1438 to S1569. Local crosslinks form within PP3, but more strikingly, PP3 crosslinks to both PP2 and PP1b, indicating interactions between the far C-terminus and the N-terminal HR. PP3 includes the only part of the PSC-CTR predicted to be structured (Fig. 1C) and overlaps a region previously implicated in interacting with Cyclin B[30]. Both secondary structure predictions and *ab initio* structure modeling (sFig. 5) [31, 32] suggest this region can form alpha helical structures. The crosslinked residues fall outside the predicted helical regions, but would bring the helical region close to the PSC HR and thus the core of PRC1.

We conclude that upon DNA binding, the conformations of the PSC-CTR are likely to be more compact, reflecting both local interactions and large looping interactions. The increased number of intra-protein crosslinks is also consistent with more dynamic behavior (sampling of more conformations) in the presence of DNA.

### Inter-protein crosslinks between PSC and dRING

We also detected inter-protein crosslinks between PSC and dRING (sFig. 2, sFig. 6). In the absence of DNA, dRING crosslinks to PP1b, the PSC RAWUL, and PP3 (sFig. 6A, sTable 3). In the presence of DNA, dRING crosslinks to PP1a and PP1b (sFig.6A, sTable 4). Intra-protein crosslinks in dRING also change on DNA binding (sFig. 6B, C, sTables 5, 6). Thus, the PSC-CTR interacts with the core of PRC1, both by interacting with the PSC-HR (PP1a), and with dRING. We did not identify high confidence crosslinks involving Pc.

### Acetylation-based protein footprinting to identify accessible lysine residues in PRC1ΔPh

XL-MS data indicate that the conformations of the PSC-CTR change on DNA binding, but do not indicate where on the PSC-CTR DNA may bind. As a means to identify DNA binding regions of the PSC-CTR, and candidate residues that contact DNA in the context of PRC1, we modified a previously described chemical acetylation based protein footprinting method[33]. In this assay, accessible lysines are chemically acetylated by treatment with NHS-acetate. Previously, NHS-acetate was used to modify accessible lysine residues, followed by MS to detect the modified sites[33]. However, in preliminary assays, we realized that it was difficult to normalize the acetylation signal across samples. This problem was exacerbated by the large number of lysine residues in PRC1 (137 in PSC, 197 in PRC1ΔPh). We therefore developed a two-step protocol (based on[34]) in which NHS-acetate was first used to acetylate accessible lysines (K-Ac). The remaining lysines were propionylated (K-Prop) by treatment with priopionic acid after protein denaturation (Fig. 4A). Thus, at each site we measured accessibility as the ratio of K-Ac/(K-Ac +K-prop). Using this method, we obtained usable data from 100 of the 137 lysines in PSC, and 154/197 of the lysines in PRC1.

**Figure 4.**
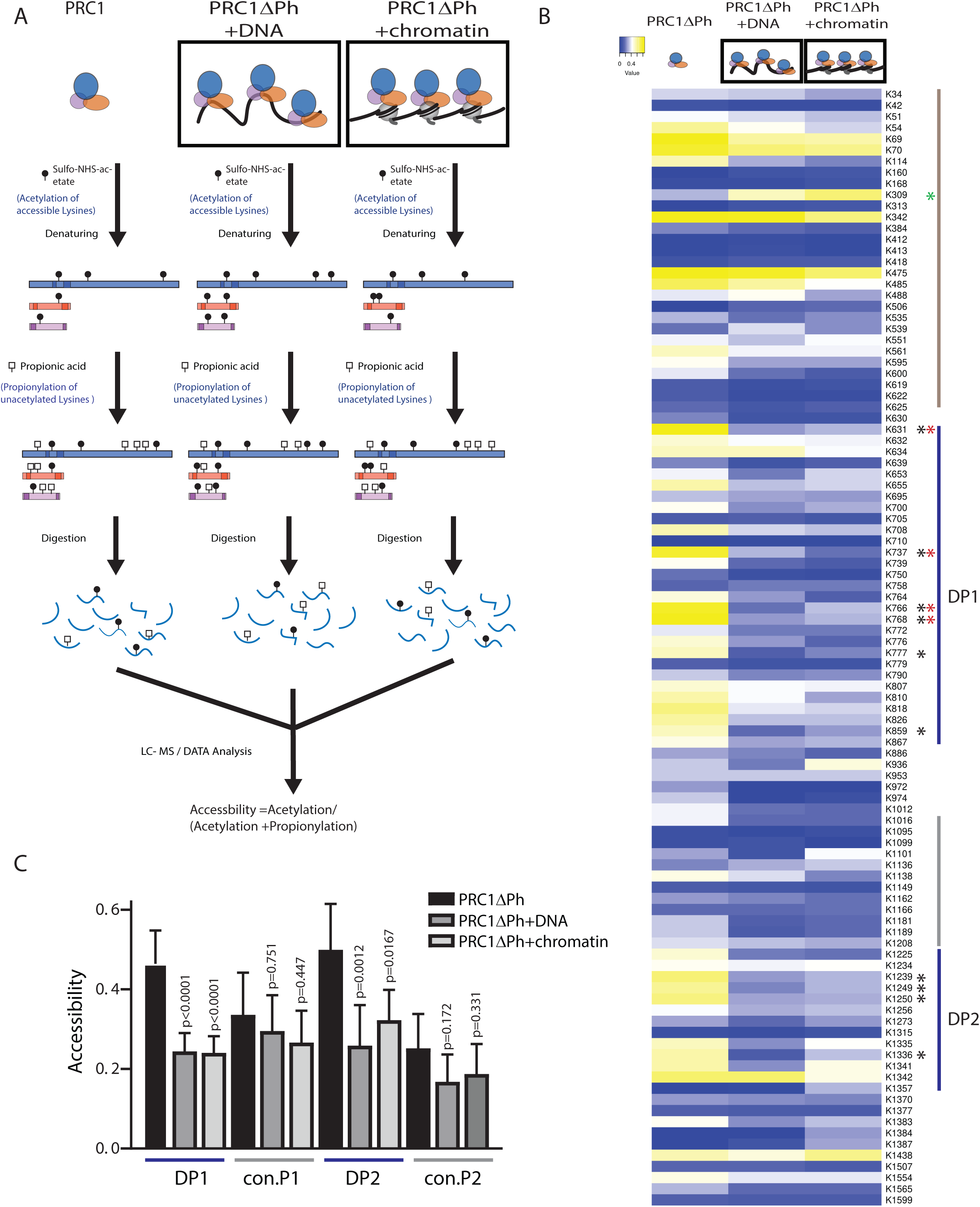
Protein footprinting by chemical acetylation of accessible lysines to identify putative DNA/chromatin binding regions in the PSC-CTR. A. Schematic of protein footprinting assay. B. Heat map of accessibility of lysines in PSC in PRC1ΔPh alone or with DNA or chromatin bound. Asterisks indicate lysines with significantly decreased acetylation in samples with DNA (black) or chromatin (red) relative to PRC1ΔPh alone. Green asterisk indicates a site that has increased accessibility when PRC1ΔPh is bound to chromatin. Blue lines indicate DNA Patch (DP) 1 and 2, and gray lines control regions used in part C. C. Average accessibility of lysines in DP1 and DP2 compared with the same number of lysines from adjacent control regions (con.P1, con.P2). Bars show average + SEM. p-values are for unpaired, two-tailed t-tests.

Lysines that are highly acetylated in this type of protocol have previously been suggested to be over-represented in lysine-based XL-MS[35]. We therefore asked if the lysines in PSC that formed crosslinks are highly accessible in the acetylation footprinting assay. We find that two prominent crosslinking sites (K342, PP1a and K475, in PP1b) are highly accessible in all samples in the footprinting assay (sFig. 7A). However, accessibility of crosslinked lysines spans the full range of detected accessibilities, so that accessibility does not predict participation in crosslinks.

### An extended, bipartite putative DNA-interacting region identified in the PSC-CTR

We compared acetylation of PSC in samples of PRC1 alone, with DNA or with the same DNA assembled into chromatin. We compiled the results from 4-5 replicates of each sample, and displayed the data as a heat map (Fig. 4B). Globally, accessibility is decreased in samples with DNA or chromatin (sFig. 7B). At the level of individual lysines many residues in PSC show little change across the samples, and these tend to be lysines that have low accessibility in PRC1ΔPh alone samples. Two patches (K631-K867 and K1225-K1342) have a pattern where most lysines are highly accessible in PRC1ΔPh alone and less accessible upon binding to DNA or chromatin. These regions are therefore candidate DNA/chromatin interacting regions. To determine which lysines most consistently change accessibility, we carried out t-tests at each site. With a 5% FDR after correcting for multiple comparisons, a small number of clustered sites are significantly changed; these patches are enlarged by using a 20% FDR (asterisks in Fig. 4B). Based on the significantly changed sites, we refer to the region from K631-K810 as DNA Patch 1 (DP1), and that from K1255-K1347 as DP2. We compared the average accessibility across DP1 and DP2 to that of an equal number of lysines from adjacent regions (Fig. 4C), which confirms the global pattern in these regions of high accessibility in the absence of DNA/chromatin, and low accessibility when DNA or chromatin are bound.

### Mutation of residues with reduced accessibility in the presence of DNA reduces the binding affinity of PRC1

As a proof of principle that residues identified by acetylation footprinting are important for DNA binding, we created a mutant version of PSC with several K-->A mutations in DP1. We mutated K737, K739, K764, K766, K768, K859 that are significantly changed, and K772, K781, and K867, which have reduced accessibility that does not reach statistical significance. We prepared PRC1ΔPh with the PSCK-->A mutations and measured DNA binding using filter binding. The mutations, which are expected to partially disrupt DP1, cause a small but significant decrease in DNA binding affinity (Fig. 5). This is consistent with these lysine residues contributing to DNA binding, but the small affect raises the possibility that they do not directly contact DNA, or that other lysines can compensate for their absence.

**Figure 5.**
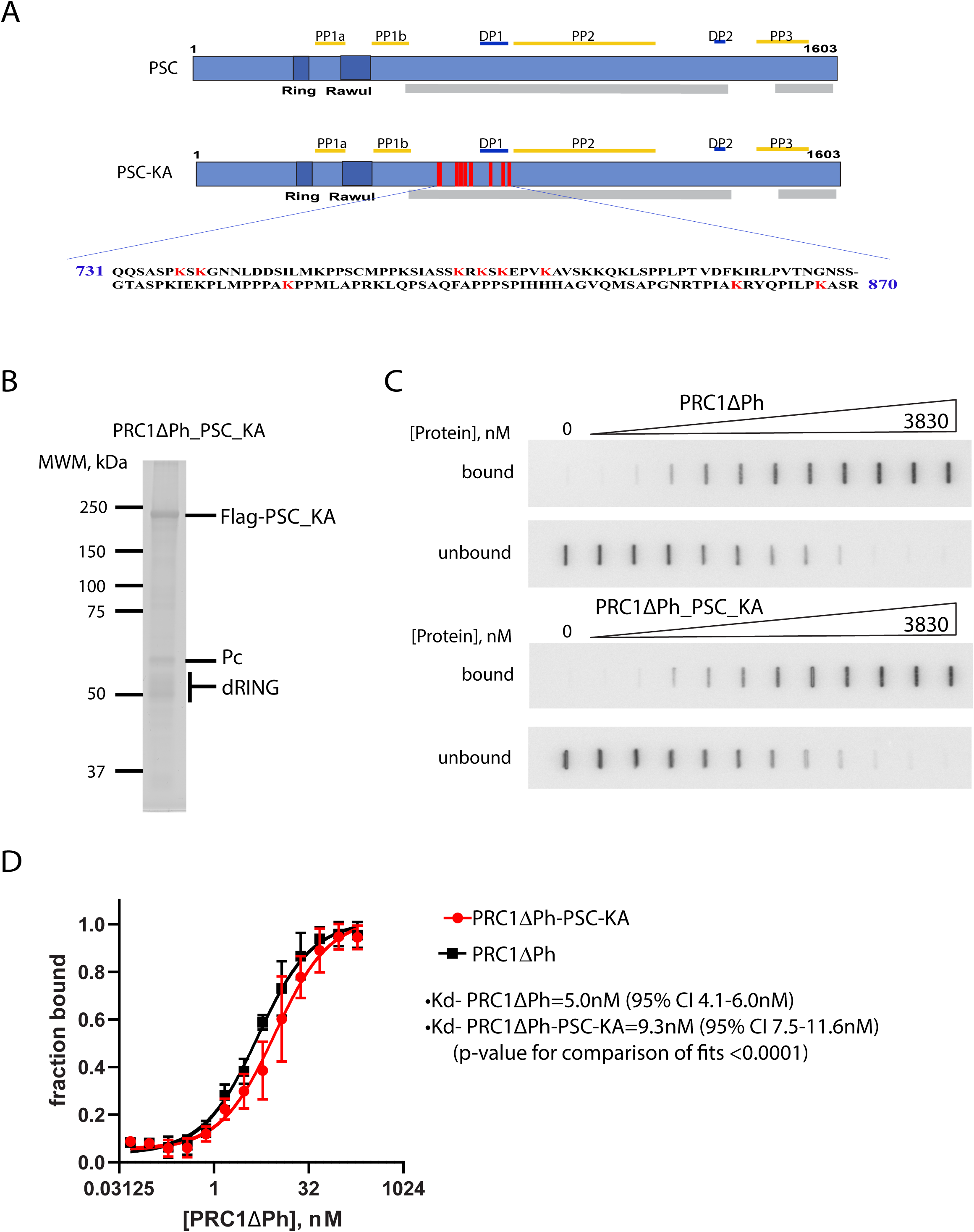
Mutation of a subset of lysines identified by protein footprinting decreases the DNA binding affinity of PRC1ΔPh. A. Schematic of PSC-KA indicating sites of K-->A mutations. B. SYPRO Ruby stained SDS-PAGE of PRC1ΔPh-PSC-KA indicating that the mutations do not affect complex formation. C. Representative membranes from filter binding assay. D. Summary of filter binding experiments using two independent preparations of PRC1ΔPh and PRC1ΔPh-PSC-KA (n=3 binding assays). Curve fits were done using the mean of the replicates using a least squares non-linear regression (Y=ABmax*X/(X+Kd)+b).

### PRC1 subunits Pc and dRING have few changes in lysine accessibility when DNA or chromatin is bound

We also analyzed lysines in Pc and dRING (sFig. 8) using the same criteria as for PSC. Only three lysines, K92 and K144 of dRING, and K170 of Pc have significantly decreased accessibility on binding DNA and/or chromatin, and no global trend is visible in heat maps for these two subunits. This is consistent with the PSC-CTR being responsible for high affinity DNA/chromatin binding by PRC1.

### Changes in lysine accessibility in histone proteins on binding PRC1ΔPh

We analyzed accessibility of histones in chromatin alone versus chromatin plus PRC1ΔPh samples. We only obtained data from 33/57 lysines, with poor representation of H2A (two sites). The data are highly variable (sFig. 9) so that no significantly changed lysines upon binding of PRC1ΔPh were identified after correction for multiple sampling. We expect that technical modifications, including use of different digestive enzymes and separation of histones from PRC1 prior to analysis could improve the analysis of histones in this assay.

Despite the limitations in the data obtained from histones, a trend towards *increased* accessibility of H3K18, H3K79, H3K122, and the tail of H4 (H4K5, H4K8, H4K12, H4K16) in the presence of PRC1ΔPh is observed (sFig. 9A). This prompted us to examine the structure of the nucleosome bound to the E3 ligase motif of PRC1, particularly since BMI1 (the PSC homologue) was previously noted to contact H3D77 in this structure[36, 37]. We find that H3K79, and the exit point of the H4 tail (the lysines in the H4 tail are not visible in the structure) are both close to K309 of PSC (sFig. 10), which is the only lysine whose accessibility is *increased* when PRC1ΔPh binds to chromatin (Fig. 4B). These data suggest a patch consisting of the tail of H4, the region of H3 around K79, and the loop containing K309 of PSC may undergo conformational alterations when PRC1ΔPh binds chromatin. H3K79 is a well-studied site of methylation that is implicated in chromatin regulation, including by the Trithorax Group (TrxG) proteins that antagonize PcG proteins[38]. Although H3K79 methylation is most commonly (but not exclusively) associated with transcriptional activation, in *Drosophila*, the *grappa* gene encoding the H3K79 methyltransferase behaves as a PcG gene (i.e. has a repressive function) in some assays and a TrxG in others, and was identified as a dominant suppressor of PcG-dependent pairing sensitive silencing[39]. Our data raise the possibility that these genetic interactions could reflect biochemical interaction between PRC1 and this region of the nucleosome, a possibility that will be interesting to explore in the future.

### A model for the PSC-CTR

To understand how the protein-protein contacts identified by XL-MS relate to DNA binding by the PSC-CTR, we compared the patches of lysines that are protected on acetylation (DP1 & DP2) to the crosslinking patches (PP1-3) (Fig. 6A). This shows that the PP and DP regions are interspersed. Only two sites (K114, K1250) are both detected in crosslinks and have decreased accessibility on binding DNA. No other lysines in the first 300 aa of PSC have altered accessibility when DNA is bound, so that the change in K114 accessibility may reflect protein-protein interaction or conformational change. In the case of K1250, both K1250 and nearby K1256 engage in crosslinks with PP1b, both with and without DNA. Because additional residues near K1250 (K1225, K1239, K1249) have decreased accessibility upon binding DNA, we consider this region part of DP2. It is possible that it can simultaneously contact DNA and interact with PP1b. We note that although the lysines that change accessibility on DNA or chromatin binding identify candidate regions of DNA/chromatin contact, we cannot formally rule out that changes in accessibility are due to protein-protein interactions within PSC. However, because we do not observe a relationship between accessibility and crosslinking (sFig. 7A), the data are consistent with (largely) distinct protein-protein and protein-DNA interacting regions.

**Figure 6.**
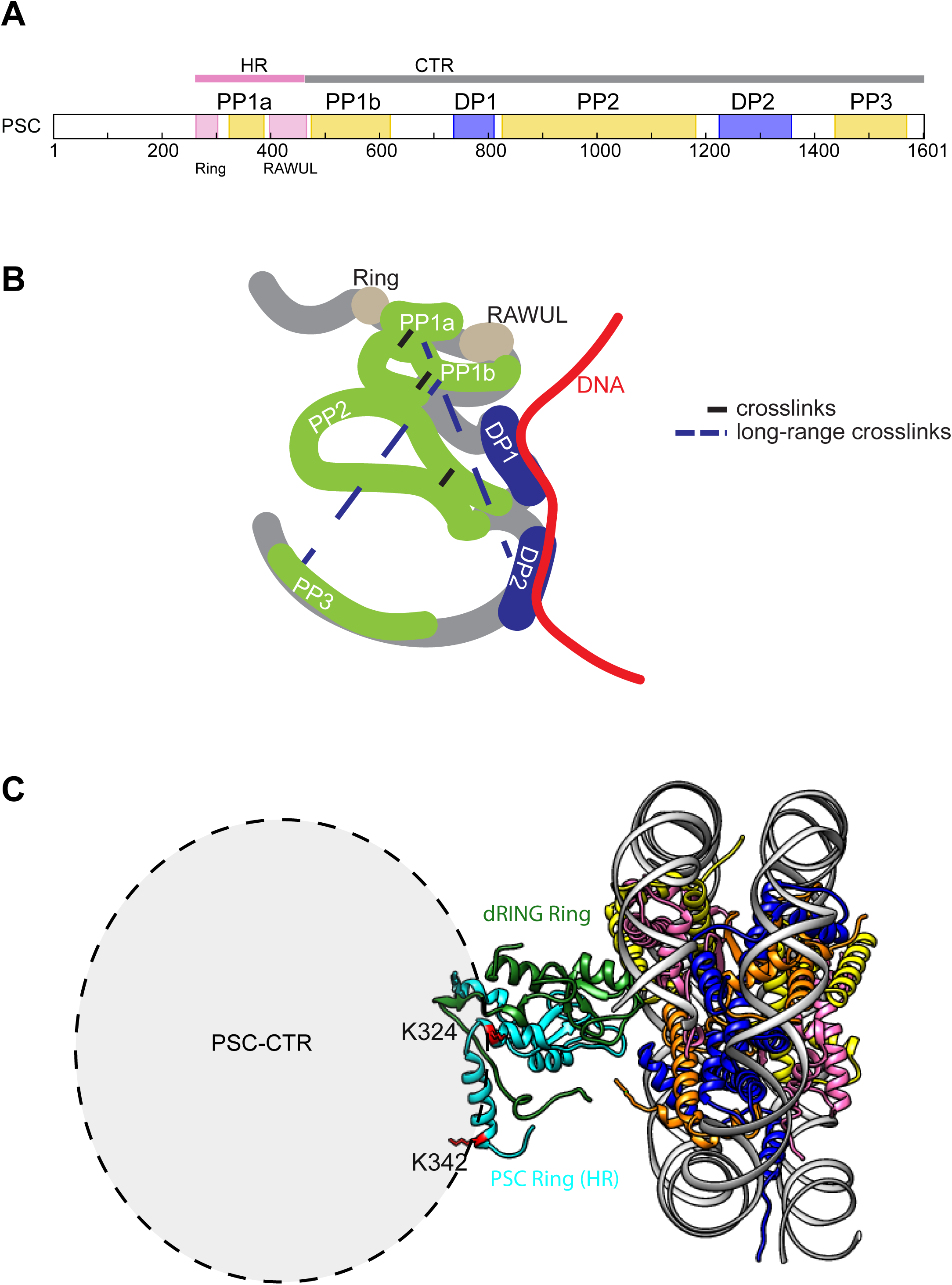
Model for organization of the PSC-CTR. A. Intra-protein interaction regions (PP) and candidate DNA/chromatin binding surfaces (DP) are largely distinct. B. Conceptual model of PSC organization. Diagram is not precisely scaled, and is meant to indicate possible folding of PSC that is consistent with crosslinking and footprinting data. Long-range crosslinks are shown by dashed lines because it was too difficult to draw these regions apposed. The model is conceptual, and not meant to imply that all contacts detected by XL-MS occur simultaneously. C. Model of the Ring domains of PSC/dRING bound to a nucleosome (based on PDB 4r8p) indicating the position of lysines in PP1A that crosslink to the PSC-CTR. Although speculative, the positions of these residues suggest the PSC-CTR (and possibly DNA or chromatin bound to it) can be brought close to the core of PRC1 without disrupting the interaction of the E3 ligase motif with the nucleosome. H3=blue, H4=orange, H2A=yellow, H2B=magenta, PSC=Cyan, dRING=green.

We previously identified aa456-909 of PSC, which include DP1, as sufficient to bind DNA[14]. The DNA binding affinity of this fragment is reduced by ∼10-fold compared with the full PSC-CTR, and it is defective in assays for nucleosome bridging, persistence through DNA replication, and inhibition of chromatin remodeling[13, 14]. Unpublished data indicate that PSC456-1350, which includes both DP1 and DP2 (separated by PP2), mirrors the activity of the full PSC-CTR (PSC456-1603). We suggest that DP1 and DP2 function as a bipartite DNA binding surface, where DNA (or chromatin) can make extended contacts. Cooperation between the two regions may be mediated by the folding of PP2 (Fig. 6B). This extended, flexible DNA/chromatin binding surface could underlay both high affinity binding, and more complex activities like nucleosome bridging and persistence through DNA replication, which may require both high affinity and dynamic DNA binding[13]. Further mutagenesis of DP1 and DP2, guided by the footprinting analysis, and of PP2 guided by the XL-MS, will be needed to test this model.

Examination of the model of the Ring fingers of PSC/dRING bound to the nucleosome shows that K324 and K342, which form crosslinks between the PSC HR and the PSC-CTR, face away from the nucleosome (Fig. 6C). Thus, although we cannot form a precise model of the PSC-CTR, these data suggest contacts between the PSC-CTR and the core of PRC1 containing the PSC HR, would be compatible with the interaction of the E3 ligase motif with the nucleosome. This may allow PRC1ΔPh to bind at least two nucleosomes simultaneously (i.e. one or more to the extended PSC-CTR surface and one to the PRC1 core), and also to ubiquitylate nucleosomes even when chromatin is compacted.

Our functional designation of DNA binding and protein folding regions of the PSC-CTR is also consistent with a set of *Psc* truncation alleles which form a phenotypic series, with progressive truncation of the PSC-CTR producing increasingly severe phenotypes[40]. Severe *Psc* alleles lack the CTR entirely, and the proteins encoded by them are not active in biochemical assays. An allele truncated partway through DP1 has severe phenotypic defects, and is partially functional *in vitro*[9]. However, *Psc* alleles that are truncated before DP2 but contain all of DP1 are viable. *Psc* alleles encoding longer proteins that retain part of PP2 have less severe phenotypes[9, 40]. This is consistent with both DP1 and DP2 being required for full activity, but DP1 being able to function (at least partially) independently. The flexible organization of the PSC-CTR and potentially long path of DNA on the protein are also consistent with a previous analysis of PRC1 binding to DNA[41]. In these experiments, the sequence specific DNA-binding protein PHO was used to recruit PRC1 to specific sites in a 4100 base pair DNA fragment. A PRC1-DNA complex was observed that contained 4 copies of PRC1. DNaseI footprinting, topology analysis, and scanning force microscopy suggested the DNA was wrapped around PRC1[41].

The combination of XL-MS and protein footprinting may be of general use for analyzing large IDRs that bind nucleic acids as well as how IDRs interact with structured regions. The protein footprinting assay may be especially useful for IDR-nucleic acid interactions. These could include protein-RNA interactions, which can also be mediated by lysine-rich IDRs[42]. In principle, other amino acids involved in nucleic acid interactions could be analyze using similar footprinting approaches[43].

## Materials & Methods

### Plasmids

Plasmids for Baculovirus expression of PRC1 were described previously[44, 45]. To create the PSC K-->A mutant, a gene block was synthesized containing the mutations (Integrated DNA Technologies). The gateway donor plasmid pDONR-PSC was digested with BamHI and BstZ171. The vector band was gel-purified, and ligated with the gene block digested with the same enzymes. The sequence of the replaced region was confirmed by Sanger sequencing. Gateway recombination (LR reaction) was used to create pFBFG-PSC-KA, which can be used for Baculovirus expression with two N-terminal Flag epitopes for purification.

### Protein expression and purification

#### Histones

Recombinant *Xenopus laevis* histones were expressed individually in *E. coli* and purified from inclusion bodies using sequential Q-sepharose and SP-sepharose ion exchange chromatography as described[46] (https://research.fhcrc.org/content/dam/stripe/tsukiyama/files/Protocols/expression.pdf). Octamers were assembled and purified by size exclusion chromatography as described[46, 47].

#### PRC1

PRC1ΔPh and PRC1ΔPh-PSC-KA were prepared from baculovirus-infected Sf9 cells as described[44]. Briefly, Sf9 cells were co-infected with viruses for expression of Flag-PSC, and Pc+dRING. After two days, cells were harvested and washed with PBS. All procedures were carried out at 4°C.Nuclear extracts were prepared as described[44, 48], and incubated overnight with M2 anti-Flag resin. Anti-Flag beads were serially washed with 20 column volumes (CV) each of BC300N, 600N, 1200N (20mM HEPES at pH 7.9, 0.2mM EDTA, 20% glycerol, 0.2mM PMSF, 0.5mM DTT, protease inhibitors and indicated [KCl] in mM (e.g. BC300=300mM KCl)), with 0.05% NP40 (indicated by “N”) and 10μM ZnCl_2_. Beads were incubated with a high stringency consisting of BC2000N without glycerol and with 1M urea for 15 min., and again serially washed with 10CV of BC1200N, BC600N, and BC300. To remove co-purifying Hsp70, beads were incubated with BC300 with 2mM ATP + 2mM MgCl_2_ at room temperature (RT) for 30 min with gentle rocking. Columns were returned to 4°C, washed with 20CV of BC300, and eluted in BC300 + 0.4mg/ml Flag peptide overnight. Elutions were combined and concentrated using Amicon 10kDa cut off concentrators to ∼1mg/ml. NP40 was added to 0.05%, and aliquots were flash frozen and stored at -80°C. To remove the Flag peptide before XL-MS and footprinting experiments, 100µl aliquots of protein were passed through 0.5ml Zeba Spin desalting columns (7MWCO, Thermo Fisher) equilibrated in BC300, according to the manufacturer’s instructions.

#### Chromatin assembly

Chromatin assembly was carried out using plasmid templates, as described (Seif et al., submitted). Briefly, plasmids containing 40 copies of the 5S nucleosome positioning sequence were assembled with recombinant *Xenopus* histones by salt gradient dialysis. Assembled chromatin was dialyzed into HEN buffer (10mM Hepes, pH 7.9, 0.25mM EDTA, 10mM NaCl) and stored at 4°C. Chromatin assembly was confirmed by EcoRI digestion followed by electrophoresis on a native 0.5X TBE-5% polyacrylamide gel, staining with Ethidium bromide, and imaging on a Typhoon imager. Chromatin assembly was further validated by Micrococcal nuclease (MNase) analysis. 200-250ng of chromatin were digested for 7 min at RT with decreasing amounts of MNase (Sigma). A 0.5U/µl stock was diluted 1:18, 1:54, 1:162 and 1:1486 into MNase dilution buffer (50mM Tris, pH 8.0, 10mM NaCl, 126mM CaCl_2_, 5% glycerol), and 1µl of each dilution used in a 10µl reaction. Reactions were stopped with DSB-PK (1% SDS, 50mM Tris-HCl, pH 8.0, 25% Glycerol, 100mM EDTA, 6.7µg/µl Proteinase K (Biobasic) and digested overnight at 50°C. MNase digests were electrophoresed on 1X TBE, 1.5% agarose (SeaKem) gels, stained with Ethidium bromide, and imaged on a Typhoon Imager.

#### Sample preparation for XL-MS

PRC1ΔPh (final [protein]=300ng/μl) alone or with DNA were mixed in binding buffer (60mM KCl, 14mM HEPES, pH 7.9, 0.3mM EDTA, 0.15mM PMSF, 2mM MgCl_2_, 1mM DTT) and incubated at 30°C for 30min in a Thermomixer at 1200rpm. BS^3^ (Thermofisher, #21580) was added to a final concentration of 2mM. After 2hr at RT, reactions were quenched by addition of ammonium bicarbonate (Sigma) to a final concentration of 0.1M. Crosslinking reactions were loaded on 600µl 5∼35% glycerol gradients (Beckman 344090 tubes) in BC300N buffer and centrifuged for 4hr at 55,000rpm at 4°C in a TLS-55 rotor (with Beckman 356860 adaptors). 40µl fractions were collected by pipetting from the top. Fractions were analyzed by SDS-PAGE followed by SYPRO-Ruby staining. Fractions containing crosslinked complexes were pooled and precipitated by addition of 4 sample volumes of ice cold acetone (Sigma), incubation at -20°C for 60min and microcentrifuged at maximum speed (14,000rpm) at 4°C for 60min. Proteins were resuspended in 5μL of 8M urea (Sigma) in 100mM ammonium bicarbonate for 10min at 30°C, reduced with 10mM DTT for 30min at 37°C and alkylated with 20mM iodoacetamide (Bioshop) for 20min at RT in the dark. Urea concentration was reduced to 2M by addition of 22.5µl of MS grade water (Fisher Scientific), and trypsin (Promega, #. V5111) added at a 20:1 (w/w) ratio. Digests were incubated for 6 hr at 37°C, followed by digestion with chymotrypsin (Sigma, # C6423) at 60:1 (w/w) overnight at 30°C. Samples were then acidified with a final concentration of 0.1% (v/v) formic acid (FA) (Sigma), and crosslinked peptides were enriched using MCX (mixed mode cationic exchange) as described[24]. In brief, peptides were reconstituted with 0.1% Trifluoroacetic acid (TFA) buffer, loaded onto an Oasis MCX µElution plate (Oasis, # 186001830BA), and washed serially with 0.1, 0.2 and 0.3M of ammonium acetate in 40% methanol and 0.1% TFA. Peptides were eluted with 2M of ammonium acetate in 80% methanol and 0.1% TFA, and evaporated to dryness. Two cycles of addition of water and evaporation were carried out to eliminate ammonium acetate. Prior to LC-MS/MS, protein digests were re-solubilized under agitation for 15min in 15µL of 0.2% FA. Desalting/cleanup was performed with C18 ZipTip pipette tips (Millpore, # ZTC1 8S0 96).

#### Sample preparation for acetylation protein footprinting

PRC1ΔPh (4µg) alone, or with 1µg DNA or chromatin were mixed in binding buffer (60mM KCl, 12mM HEPES, pH 7.9, 0.3mM EDTA, 0.15mM PMSF, 2mM MgCl_2_, 1mM DTT) and incubated at 30°C for 15min in a Thermomixer at 1200rpm. A freshly prepared 15mM stock of Sulfo-NHS-acetate in BCO (Thermo Fisher, # 26777) was added to a final concentration of 0.5mM, and acetylation reactions incubated at RT for 15min. TFA was added to a final concentration of 1% to quench the reactions. NaDOC (MP Biomedicals) was added to a final concentration of 0.1%, and reactions were incubated on ice for 30min. 100% TCA (Sigma) was added to a final concentration of 10%. After incubation for 16hr at 4°C, reactions were centrifuged at full speed in a microcentrifuge for 1hr at 4°C. Pellets were washed twice with 100% acetone and dried at RT. Pellets were resuspended in 8M urea/100mM NH_4_HCO_3_ and incubated for 10min at 30°C on a thermomixer (1200rpm). DTT in 100mM NH_4_HCO_3_ was added to a final concentration of 9mM and reactions incubated for 20min at 37°C and 350rpm. Reactions were equilibrated to RT for 10min and iodoacetimide (100mM in 100mM NH_4_HCO_3_) added to a final concentration of 10mM for 20min in the dark. Reactions were diluted 1:2 with H_2_O. Propionic anhydride (Sigma) was prepared with acetonitrile (ACN) (1:3, v/v) and added to acetylation reactions (1:4, v/v) for 15min at RT. Propionic anhydride was added again (1:3 v/v), and reactions incubated a further 15 min. Reactions were completely dried by speed vac, resuspended in H_2_O and propionylation repeated a third time as described above. Reactions were dried by speed vac, resuspended in 50mM NH_4_HCO_3_, and sequentially digested with trypsin and chymotrypsin as described above.

### Mass Spectrometry

#### XL-MS

Eluates were dried down in a speed vac, re-solubilized under agitation for 15min in 40 of 2%ACN-1%FA. 9.5µL were injected into a 75μm i.d. × 150mm Self-Pack C18 column installed in an Easy-nLC II system (Proxeon Biosystems). The buffers used for chromatography were 0.2% FA in water (Buffer A) and 0.2% FA in 100% ACN (Buffer B). Peptides were eluted with a four slope gradient at a flowrate of 250nL/min. Solvent B first increased from 2 to 4% in 6 min, then from 4 to 7% B in 8 min, then from 7 to 34% B in 96 min and finally from 34 to 85% B in 10 min. The HPLC system was coupled to Orbitrap Fusion mass spectrometer (Thermo Scientific) through a Nanospray Flex Ion Source. Nanospray and S-lens voltages were set to 1.3-1.7kV and 70V, respectively. Capillary temperature was set to 250 °C. Full scan MS survey spectra (m/z 360-1560) in profile mode were acquired in the Orbitrap with a resolution of 120,000 with a target value at 1e6. A top 3 second (sec) method was used to select the most intense peptide ions for fragmentation in the HCD collision cell and analysis in the Orbitrap with a target value at 5e4 and a normalized collision energy at 32. Target ions selected for fragmentation were dynamically excluded for 30sec after 2 counts.

#### Acetylation protein footprinting

Prior to LC-MS/MS, protein digests were re-solubilized under agitation for 15 min in 75µL of 2%ACN/1% FA. 4µl were injected into a 75μm i.d. × 150mm Self-Pack C18 column installed on the Easy-nLC II system (Proxeon Biosystems). Peptides were eluted with a two slope gradient at a flowrate of 250nL/min. Solvent B first increased from 2 to 35% in 105min and then from 35 to 85% B in 15min. The HPLC system was coupled to an Orbitrap Fusion mass spectrometer (Thermo Scientific) through a Nanospray Flex Ion Source. Nanospray and S-lens voltages were set to 1.3-1.7kV and 60V, respectively. The ion transfer tube temperature was set to 250°C. Full scan MS survey spectra (m/z 360-1560) in profile mode were acquired in the Orbitrap with a resolution of 120,000 and a target value at 3e5. The 25 most intense peptide ions were fragmented in the HCD collision cell and analyzed in the linear ion trap with a target value at 2e4 and a normalized collision energy at 28. Target ions selected for fragmentation were dynamically excluded for 25sec after 2 counts.

#### Analysis of XL-MS data

A customized protein data base containing the three PRC1 subunits with trypsin and chymotrypsin as digestion enzymes and up to three missed cleavages was used for all analysis. Carbamidomethylation of C was set as a fixed modification, and oxidation of M and phosphorylation of S, T, ad Y as variable modifications.

#### pLink

Raw files were searched using pLink 2.3.7.Searches were conducted in combinatorial mode with a precursor mass tolerance of 10ppm, a fragment ion mass tolerance of 20ppm, and false discovery rate of 5%. Results were filtered by applying a precursor mass accuracy of ±10ppm.

#### MeroX

Raw files were converted to the mxML format using MSConvert (3.0). Crosslinked peptides were identified using version 2.0.beta.5 of MeroX software, and mass precision tolerances of 10 and 20ppm for precursors and fragment ions, respectively.

#### SIM-XL

Single raw files were used as input with SIM-XL (1.5.4.0). The precursor mass error was set to 10ppm and the fragment ion mass error to 20ppm. The fragmentation method was HCD, with a minimum peptide mass of 600Da and a maximum of 6,000Da.

#### Analysis of Acetylation Footprinting data

Acetylation footprinting data were analyzed in MaxQuant (v16.10.43)[49] using the following parameters:

> Fixed modifications: carbamidomethylation
>
> Variable modifications: Acetyl (K), Prop (K), Oxidation(M), Acetyl (Protein N-term)
>
> Digestion: Trypsin/P, chymotrypsin +
>
> Missed cleavages: 10*
>
> Modifications per peptide: 7
>
> Match between runs: yes
>
> Precursor tolerance: 20ppm
>
> MS/MS tolerance: 20ppm

*Because acetylation or proprionylation will block cleavage next to K, we allowed a high number of missed cleavages. The majority of peptides were identified with 8 missed cleavages or less.

Intensities for each modified site were obtained from the “AcetylK” and “PropK” sites tables for each position. Only lysines with intensities for Prop or Acetyl in at least one sample were considered. To calculate “Accessibility”, the ratio was calculated for each sample at each site as follows: (K-Ac/(K-Ac+pK-Prop+0.5)). Sites that are not acetylated score as 0, while those with acetylation only score as 1. Accessibility was averaged across samples (n=4-5) to produce the heat maps shown in Fig. 3B and sFig. 7 and 8. Heat maps were generated using <u>heatmapper.ca</u>[50]; accessibility is displayed (i.e., values are not normalized across rows or columns), and no clustering was applied. To determine which sites were significantly altered, GraphPad Prism (v8.4) was used to conduct t-tests at each site between PRC1ΔPh alone and PRC1ΔPh + DNA or + chromatin. Samples were assumed to have the same SD, and the two-stage linear step-up procedure of Benjamini, Krieger, and Yekutieli [51] in GraphPad Prism, with Q=20% was used to correct for multiple comparisons.

To visualize crosslinks and acetylation sites on the structure of Bmi-1/RING1B bound to a nucleosome (PDB4r8p), SWISS-MODEL[52] was used to generate a homology model using the sequences of PSC and dRING, and UCSF Chimera (v1.14) [53] used to generate figures and measure distances. For *ab initio* structure prediction of PSC aa1401-1580, the sequence was submitted to QUARK [31, 32]. The top predicted model was visualized in Chimera and crosslinked sites indicated.

To align PSC with human PCGFs, T-coffee[54] (http://tcoffee.crg.cat/apps/tcoffee/index.html) was used with BoxShade (https://embnet.vital-it.ch/software/BOX_form.html).

#### Filter Binding

The filter binding assay was carried out as described in[45, 55]. Briefly, a gel-purified, ^32^P body-labeled PCR product (150 bp) was used at 0.02nM. Reaction conditions (20µl volume) were as follows: 12mM Hepes, pH 7.9, 60mM KCl, 0.24mM EDTA, 12% glycerol, 0.01% NP40, 1mM DTT, 2mM MgCl_2_. PRC1ΔPh was serially diluted in 2-fold steps. Binding reactions were incubated at 30°C for 45min. Top (nitrocellulose) filters were treated for 10min with 0.4M KCl, neutralized using ddH2O, and equilibrated for at least 1 hour in filter binding buffer (same as reaction conditions without NP40). Bottom (Hybond+) filters were equilibrated in 0.4M Tris, pH 8.0 for at least 5 min. After reaction incubation, filters were assembled, and each slot washed with 100µl binding buffer. Reactions were immediately loaded, followed by 2*100µl washes in binding buffer. Filters were dried and exposed to phosphorimager screens for quantification on a Typhoon imager. Quantification was done with ImageQuant. Fraction bound was calculated in Excel using background subtracted intensities for top (bound) and bottom (unbound) membranes. Fraction bound=(top/(top+bottom)). Binding constants were calculated in GraphPad Prism.

The instrument mzML file, detailed instrument settings, MeroX results file, FASTA file containing PRC1 sequences, and MeroX settings file can be downloaded at MassIVE **<u>MSV000085418</u>**. The same accession number can be used to access the raw files for the acetylation footprinting analysis, and the MaxQuant output.

## Acknowledgements

We are grateful to Drs. Francois Robert, Marlene Oeffinger, and Christian Trahan for comments on the manuscript, to Marlene Oeffinger and Christian Trahan for technical and intellectual guidance throughout the project, and to Carolina Aguilar, and all members of the mass spectrometry core facility at the IRCM for technical assistance. This work was funded by grants from the Canadian Institutes for Health Research (311557), and the NIH (GM114338-02).

## Author contributions

**Jin Joo Kang:** Conceptualization, Investigation, Writing; **Denis Faubert**: Investigation, Writing, Formal analysis; **Jonathan Boulais:** Formal analysis; **Nicole Francis:** Conceptualization, Writing, Supervision, Funding Acquisition

## Supplementary Figure captions

**sFigure 1.**
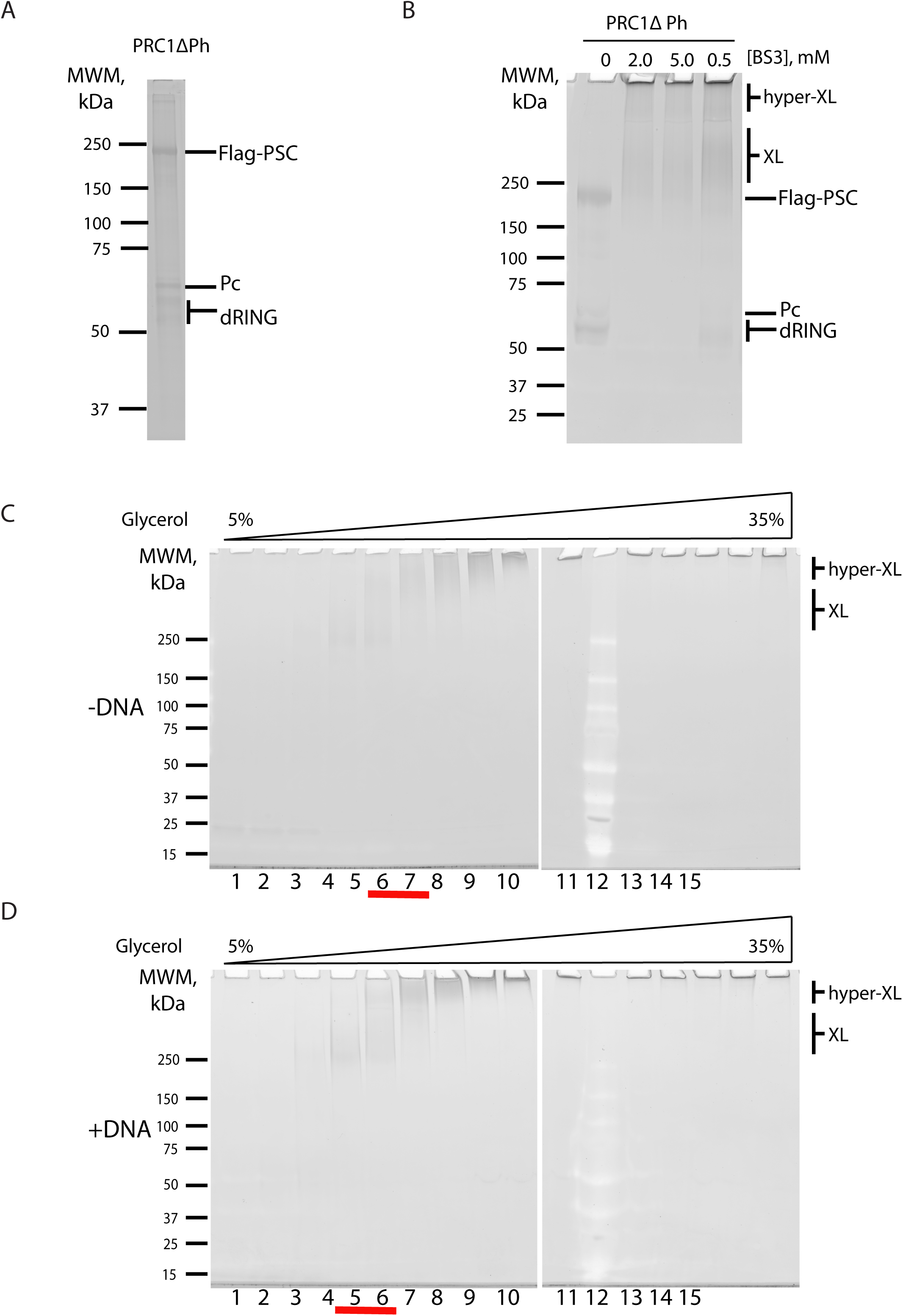
Sample preparation for XL-MS analysis of PRC1ΔPh. A. SYPRO Ruby stained SDS-PAGE of PRC1ΔPh. B. SYPRO Ruby stained SDS-PAGE of PRC1ΔPh after crosslinking with BS^3^. C. SYPRO Ruby stained SDS-PAGE of glycerol gradient fractions of crosslinked PRC1ΔPh in the absence (C) or presence (D) of DNA. Red line indicates fractions that were pooled for MS analysis.

**sFigure 2.**
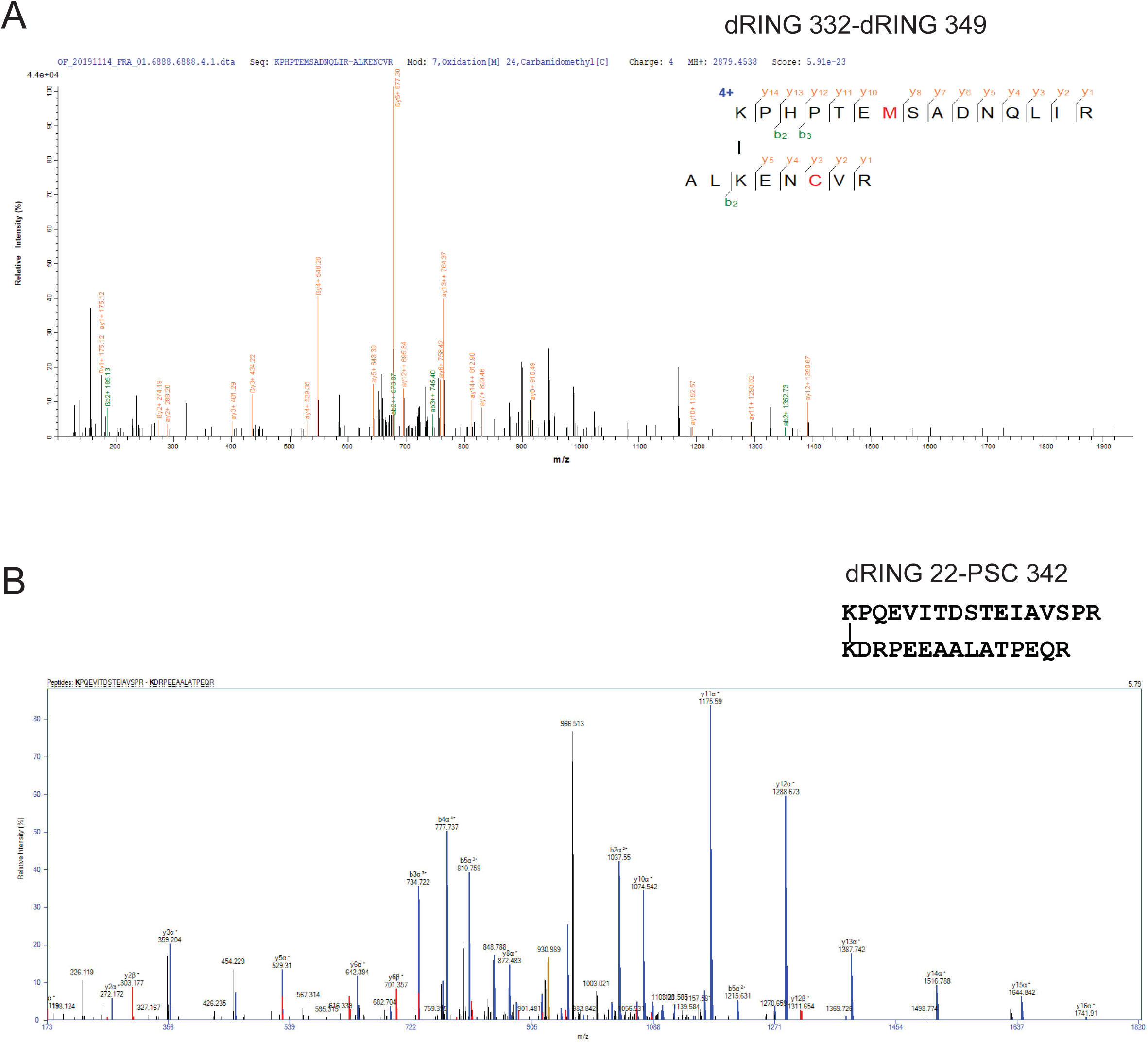
Representative spectra of crosslinked peptides identified by pLink and SIM-XL. A. Representative spectrum of crosslinked peptides identified by pLink. The dark blue, blue and red respectively correspond to a, b and y ions of peptide A. The dark green, orange and green correspond to a, b and y ions of peptide B. The red highlighted cysteine “C” is carbamido-methylated. B. Representative spectrum of crosslinked peptides identified by SIM-XL.

**sFigure 3.**
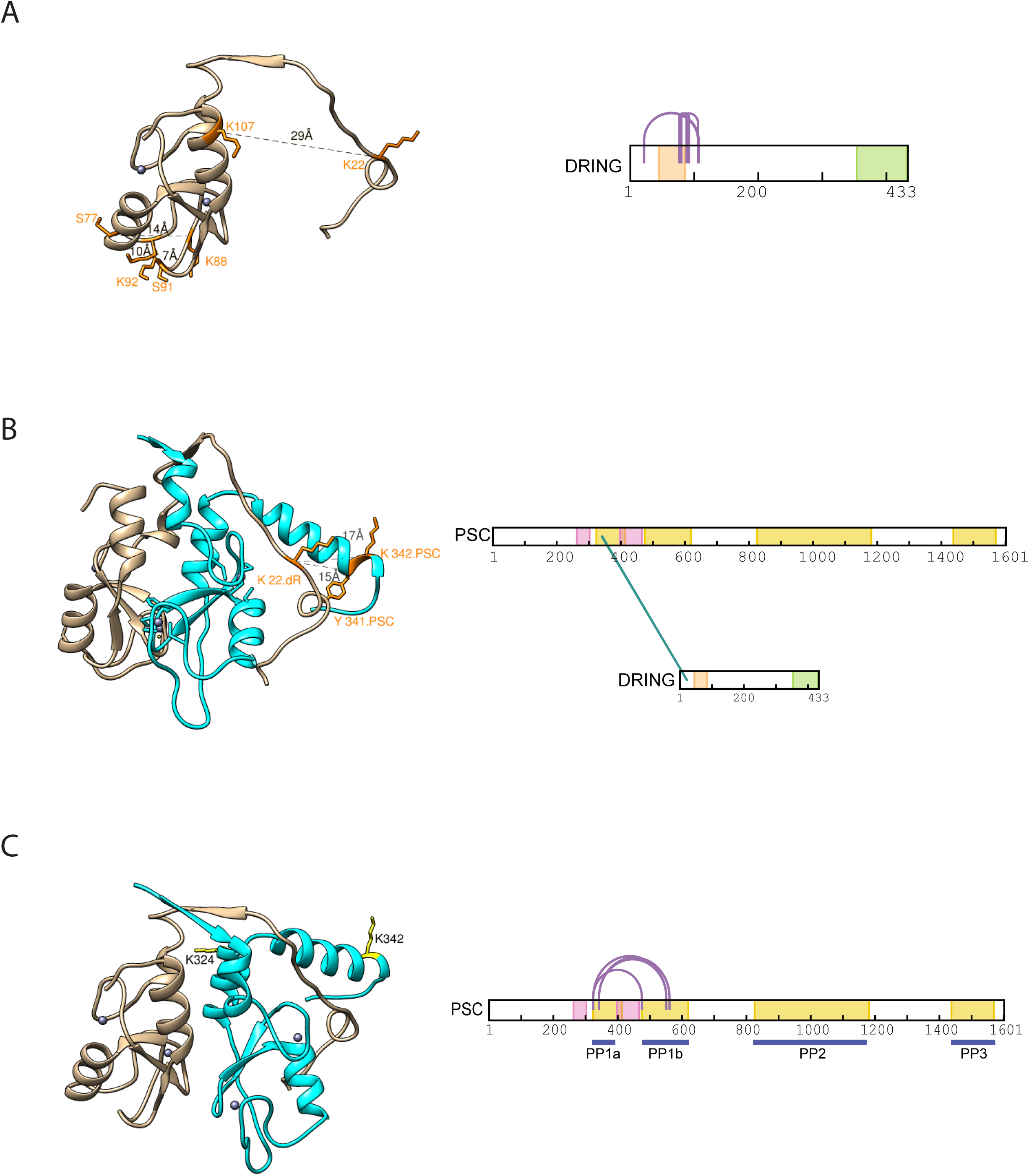
Mapping of identified crosslinks on structured domains of dRING and PSC. A. Intra-protein crosslinks in dRING modeled onto the structure of dRING in complex with PSC (PSC is not shown.) (left). Right schematic indicates positions of crosslinks. See also sTables 5, 6. B. Inter-protein crosslinks between dRING and PSC modeled onto the same structure as in A (left). PSC is in cyan, and dRING in tan. Right schematic indicates positions of crosslinks. See also sTable 4. C. Positions of crosslinked residues (K324, K342, both of which crosslink to PP1B) in PSC shown on the same structure as in A and B (left). Structural information is not available for the partner crosslinked residues. Right schematic indicates positions of crosslinks. See also sTables 1, 2. The homology model of PSC-dRING was generated using PDB 4r8p (BMI1-RING1B).

**sFigure 4.**
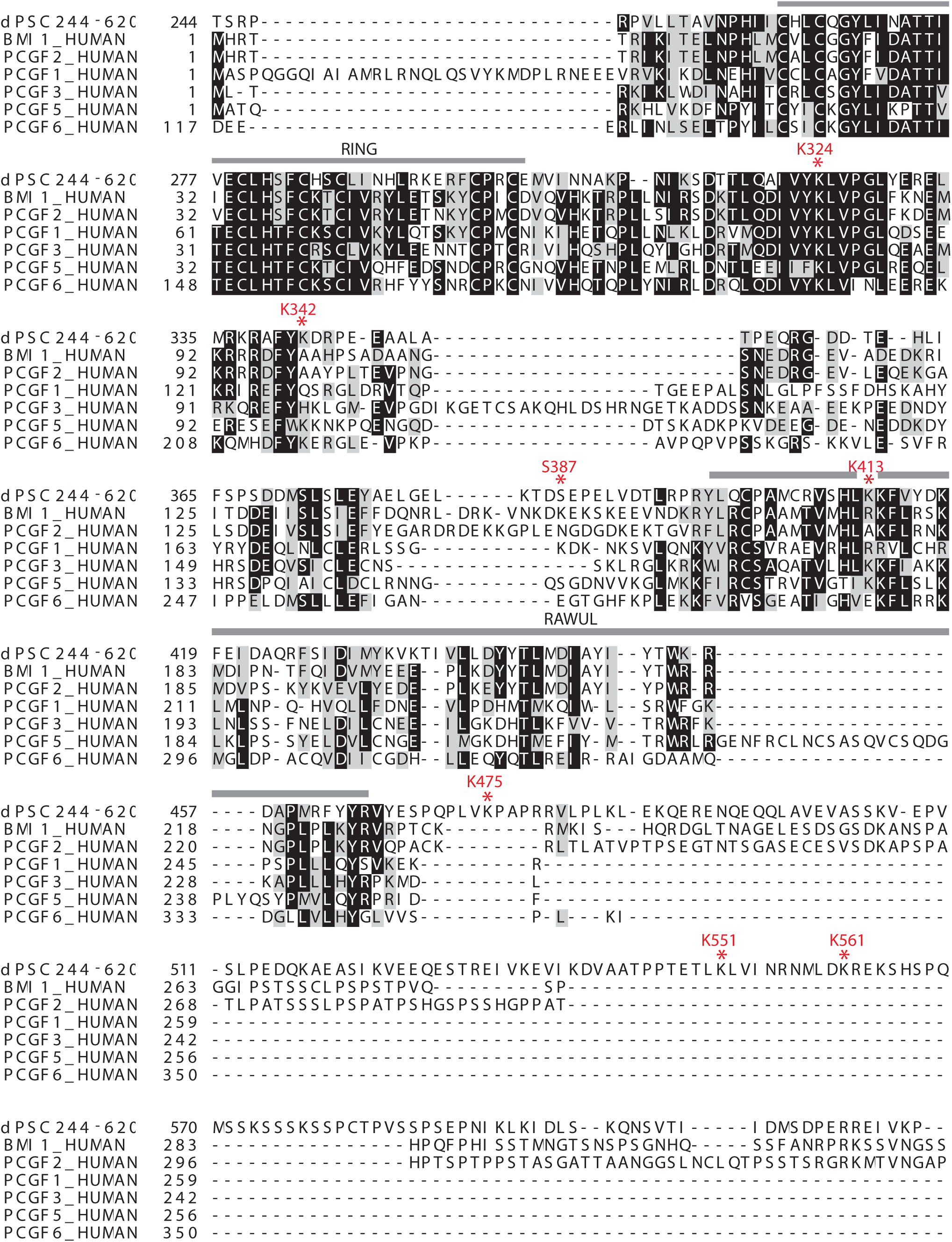
Alignment of human homologues of PSC (PCGF1-6). PSC (starting at aa244) was aligned to the 6 human homologues. BMI1 is PCGF4. Asterisks indicate residues identified in crosslinks.

**sFigure 5.**
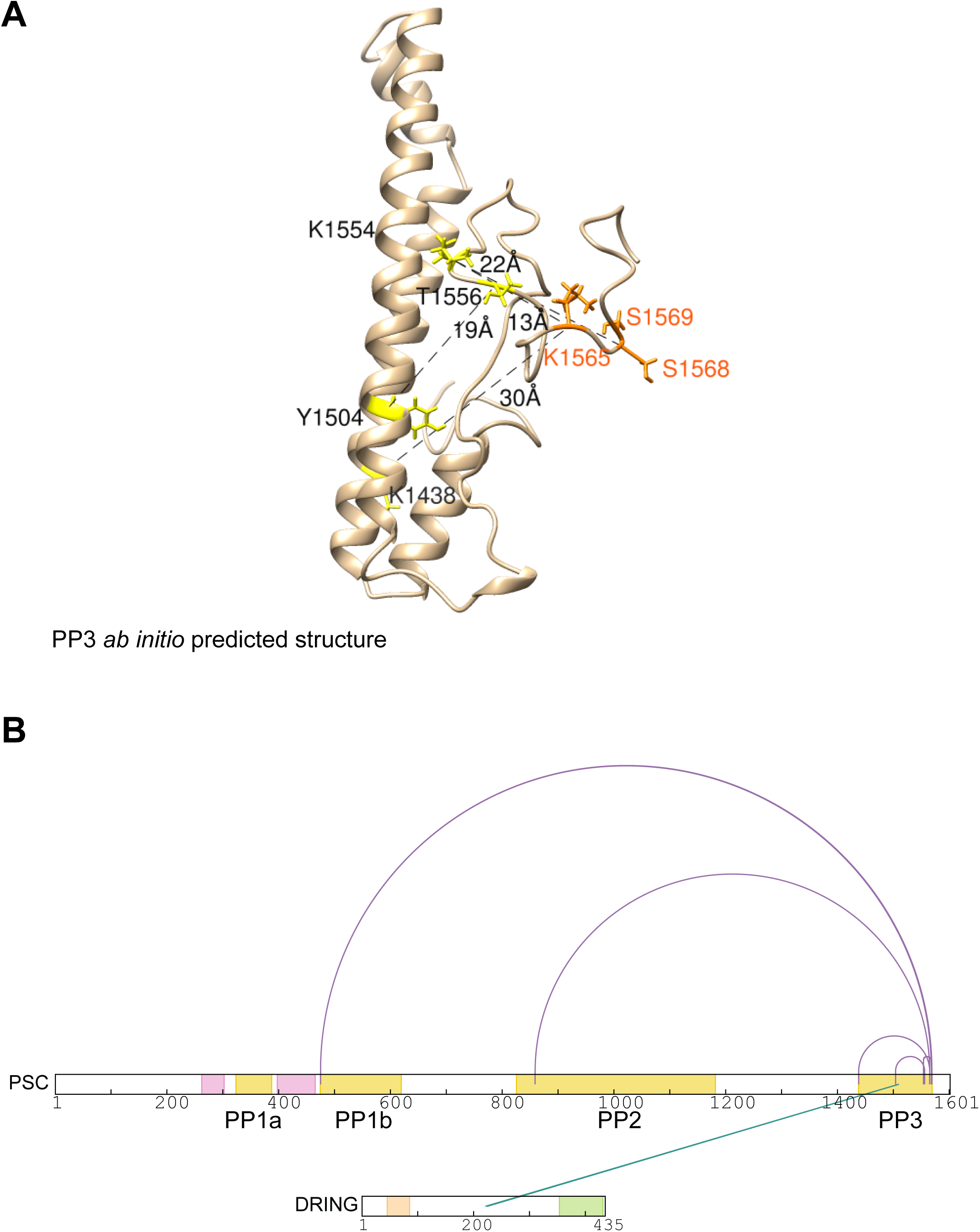
Protein Patch 3 structure prediction. A. *ab initio* predicted conformation of aa1401-1580 generated with QUARK. Crosslinks within PP3 (between yellow residues) are indicated in black. Orange residues crosslink to PP1b and PP2. B. Map of crosslinks within PP3, and from PP3 to PP2, PP1b, and dRING. See also sTable 2 for scores.

**sFigure 6.**
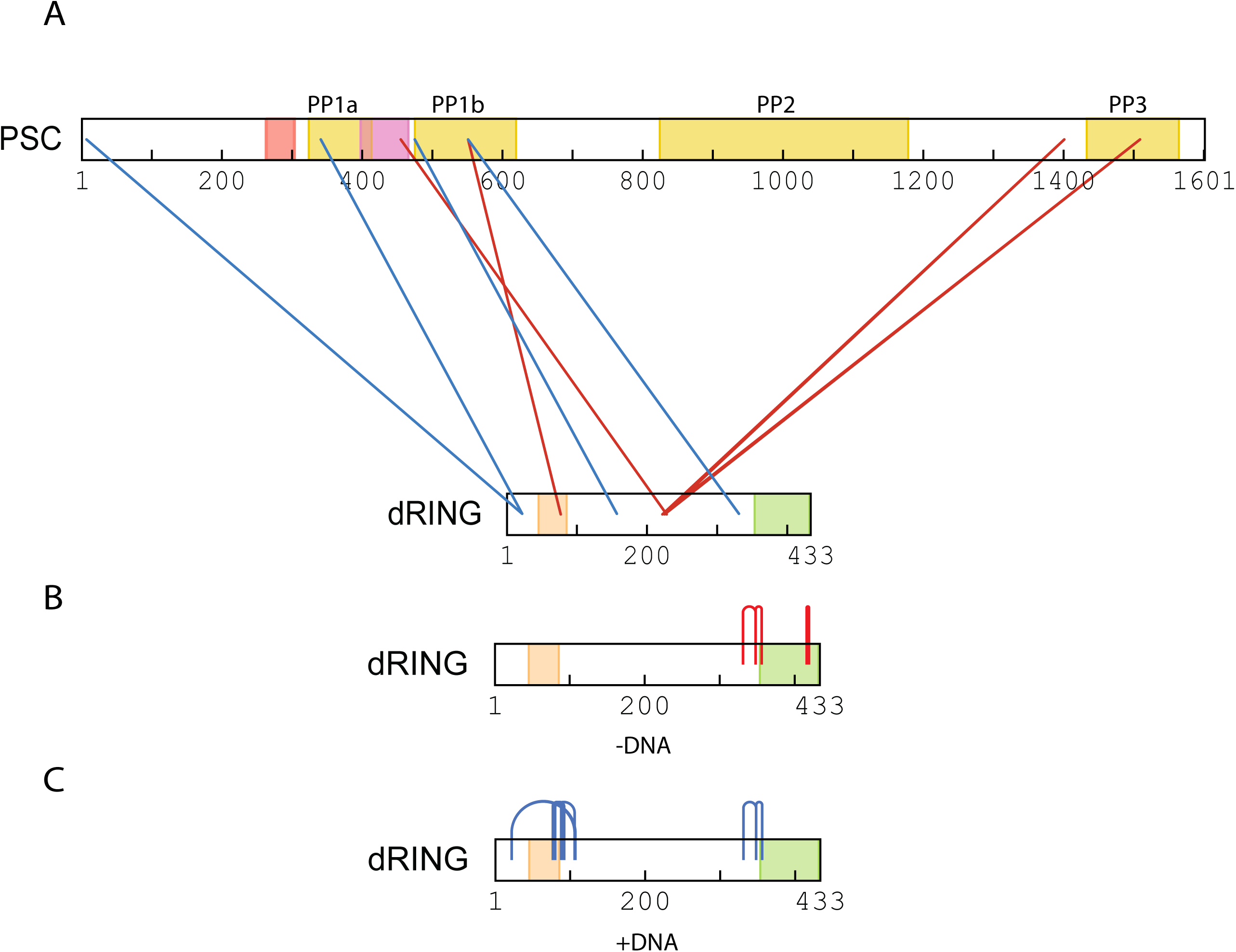
Crosslinks identified in dRING. A. Inter-protein crosslinks with PSC (red=without DNA, blue=with DNA). B, C. Intra-protein crosslinks without (B) or with (C) DNA.

**sFigure 7.**
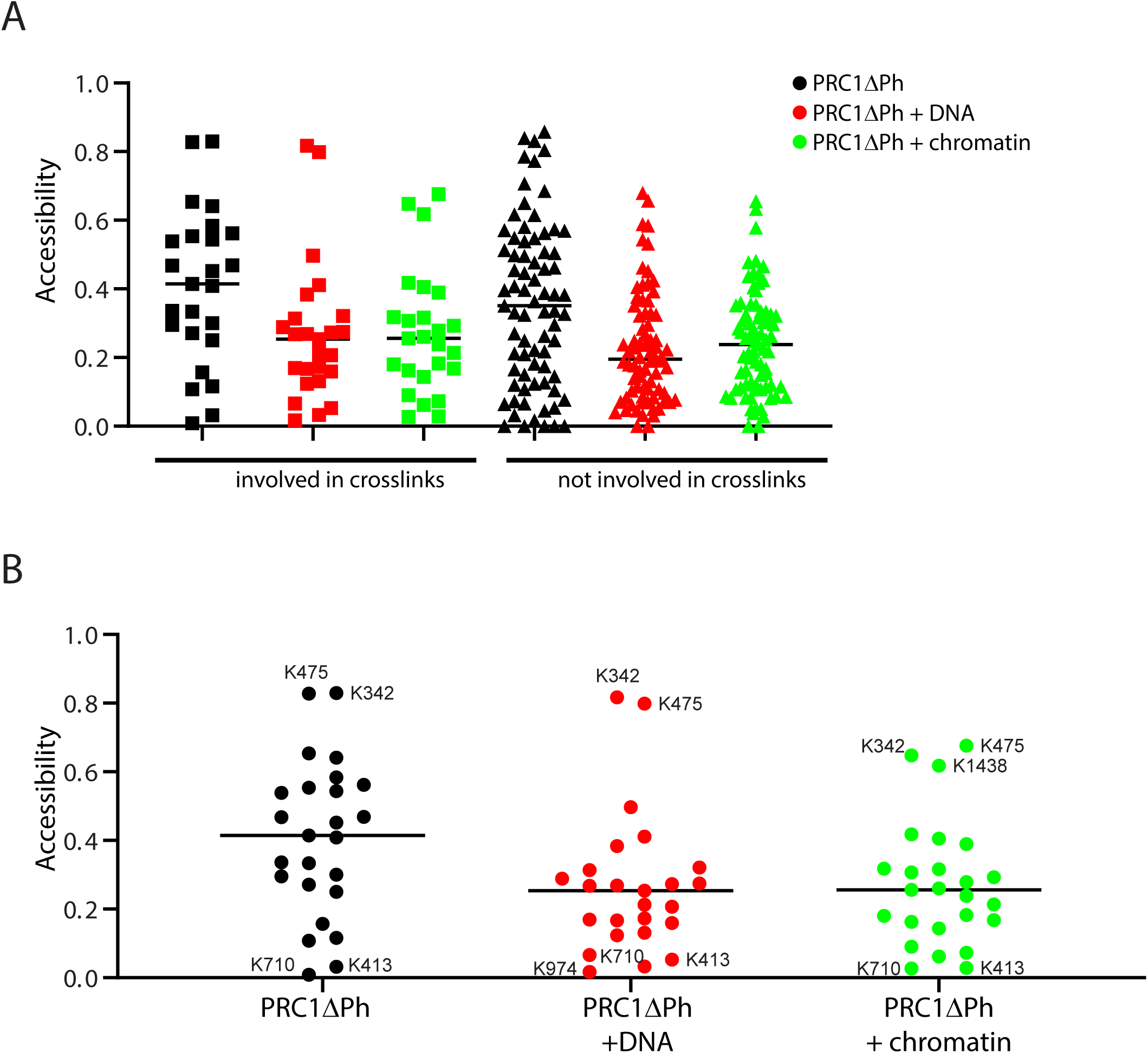
Relationship between lysine accessibility and crosslinking of candidate DNA binding patches. A. Scatter plot of accessibility of all lysines that were (left) or were not (right) identified in crosslinks. B. Scatter plot of accessibility of all lysines in the three different samples. The difference between PRC1ΔPh and PRC1ΔPh+ DNA (P<0.0001) or chromatin (p=0.0002) is statistically significant by Mann Whitney test.

**sFigure 8.**
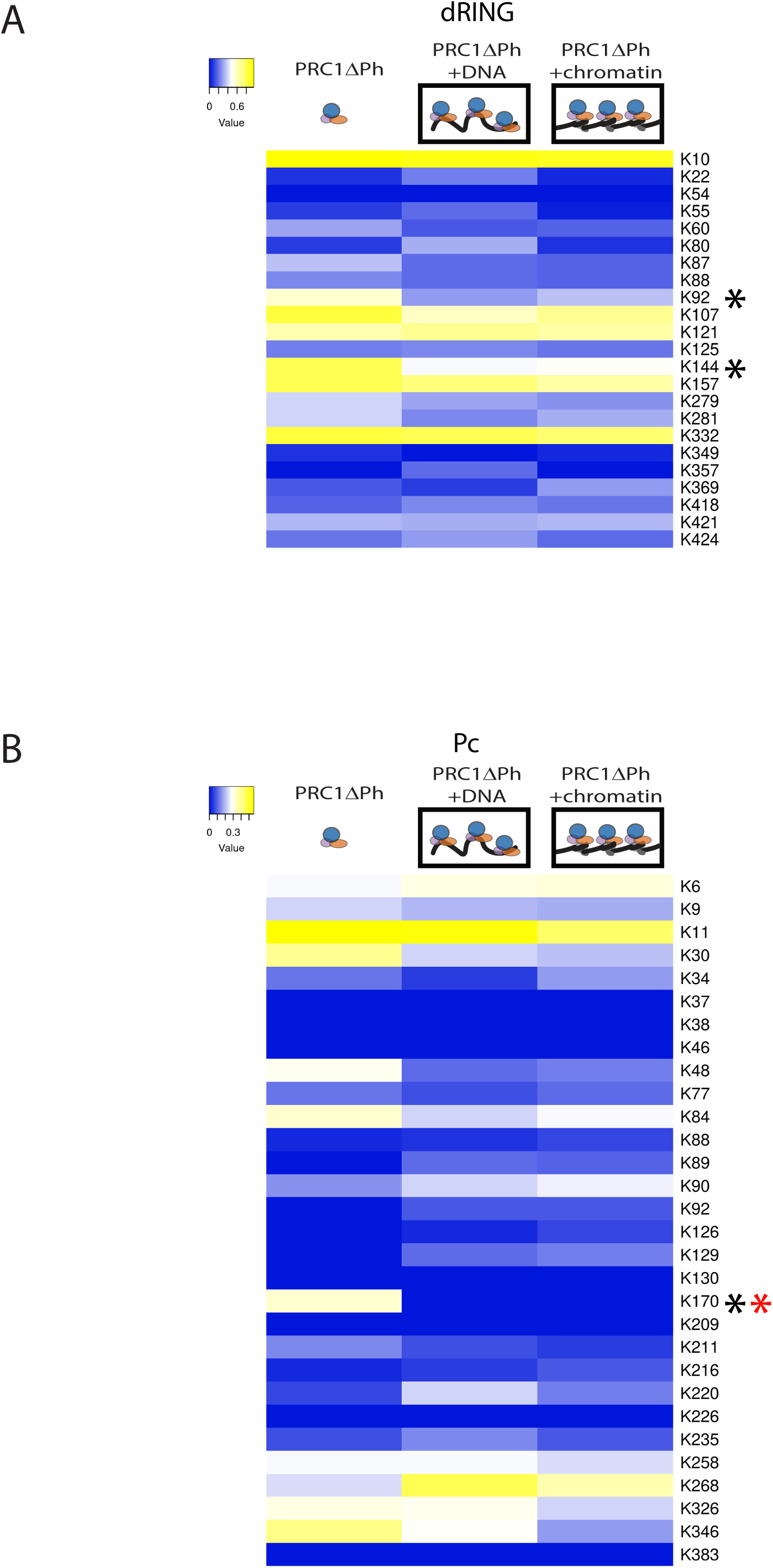
Protein footprinting of Pc and dRING. Heat maps of accessibility of dRING (A) and Pc (B). Asterisks mark lysines with decreased accessibility in the presence of DNA (black) or chromatin (red).

**sFigure 9.**
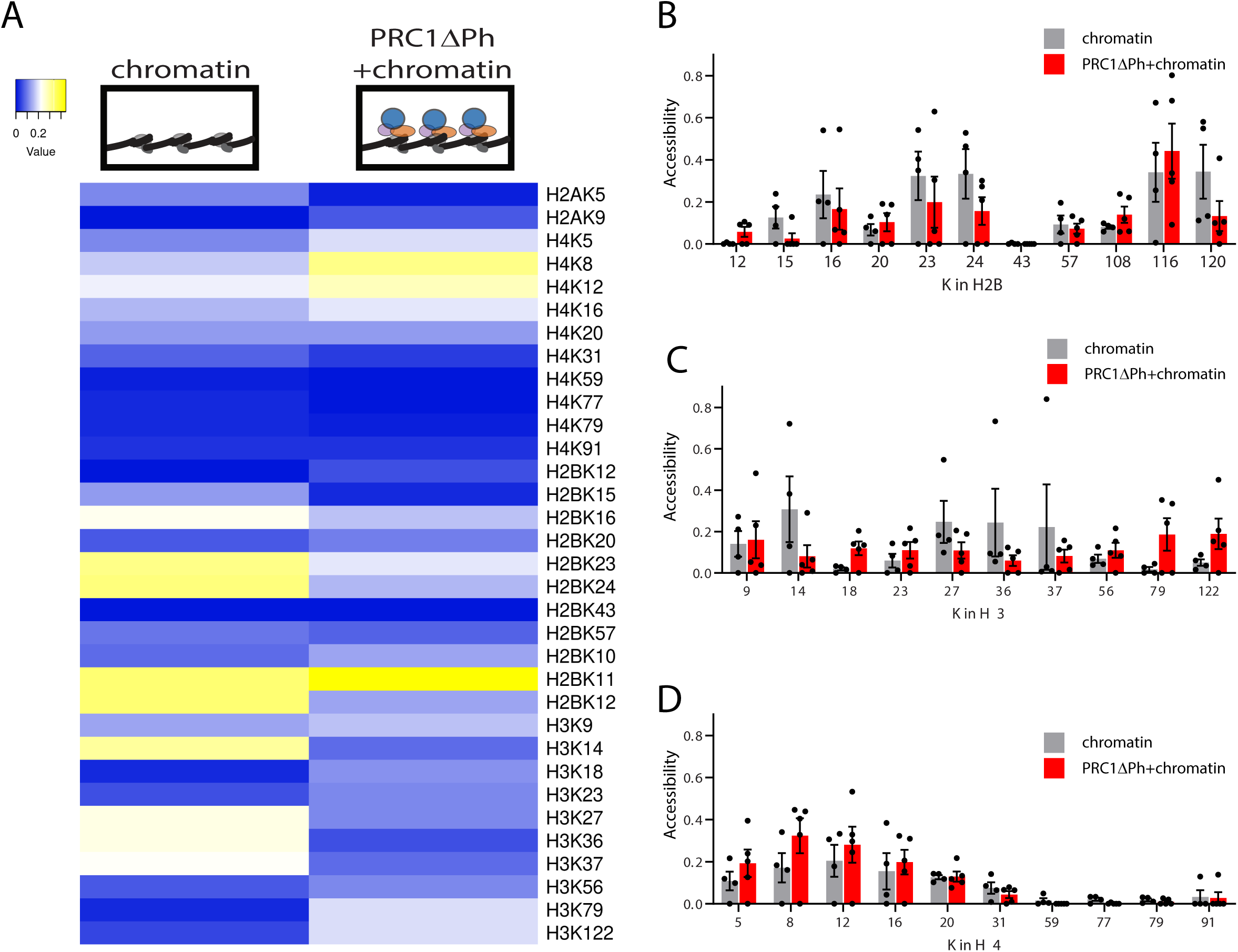
Protein footprinting of histones. A. Heat map of accessibility of histone proteins. None of the observed differences attained statistical significance. B-D. Graphs of mean +SEM of each site in histones H2B, H3, and H4 for which data were obtained. H2A is not shown because we only obtained usable data from two lysines.

**sFigure 10.**
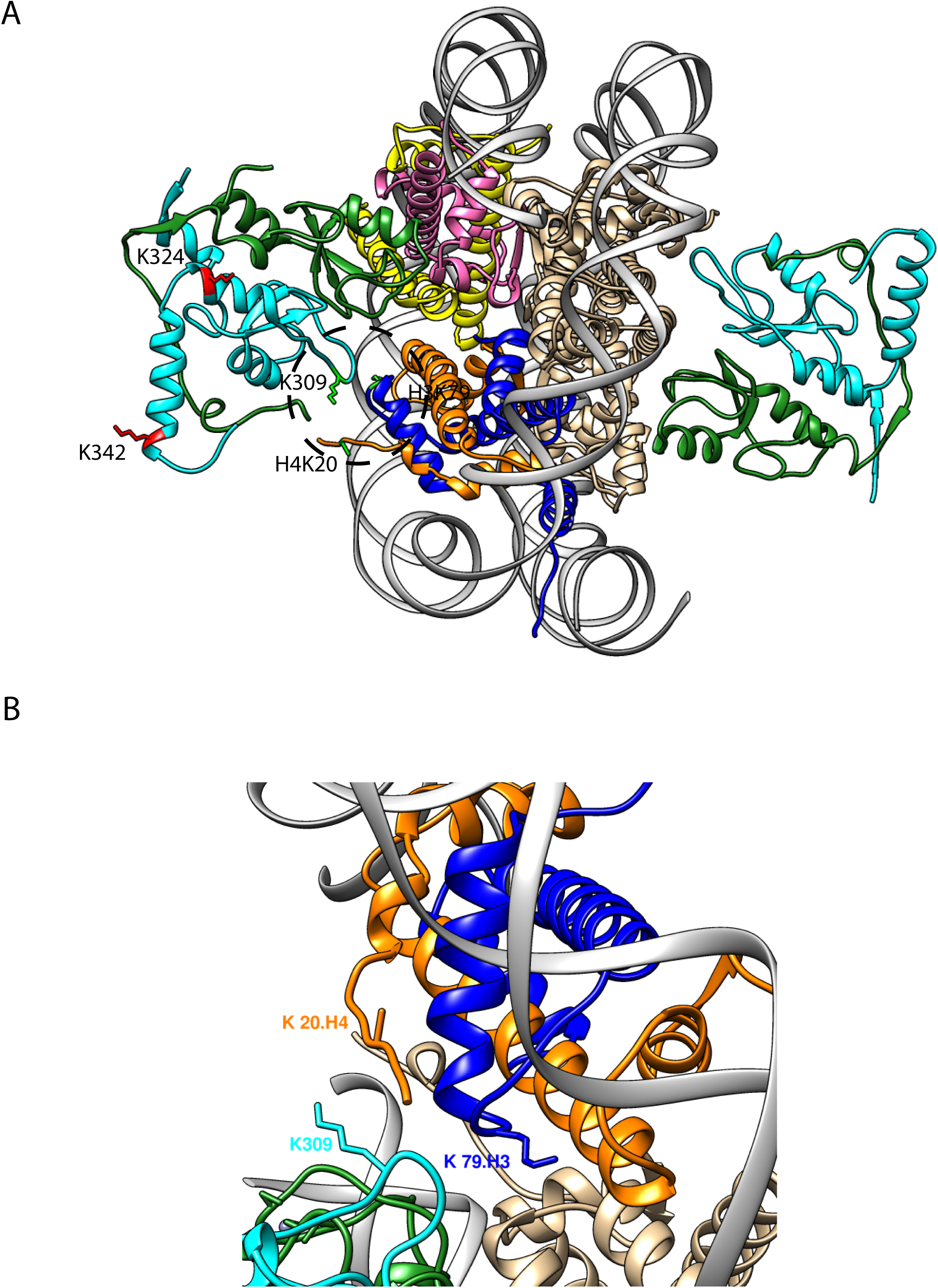
A patch of increased accessibility in PSC and the nucleosome? A Homology model of PSC (cyan)-dRING (forest green) bound to a nucleosome (based on PDB4r8p). (H3=blue, H4=orange, H2A=yellow; H2B=purple). Note that the ubiquitin E2, which is part of the structure, is not shown. H3K79, and PSCK309 are displayed in green. The H4 structure ends at residue R17.

**Supplementary Table 1.**
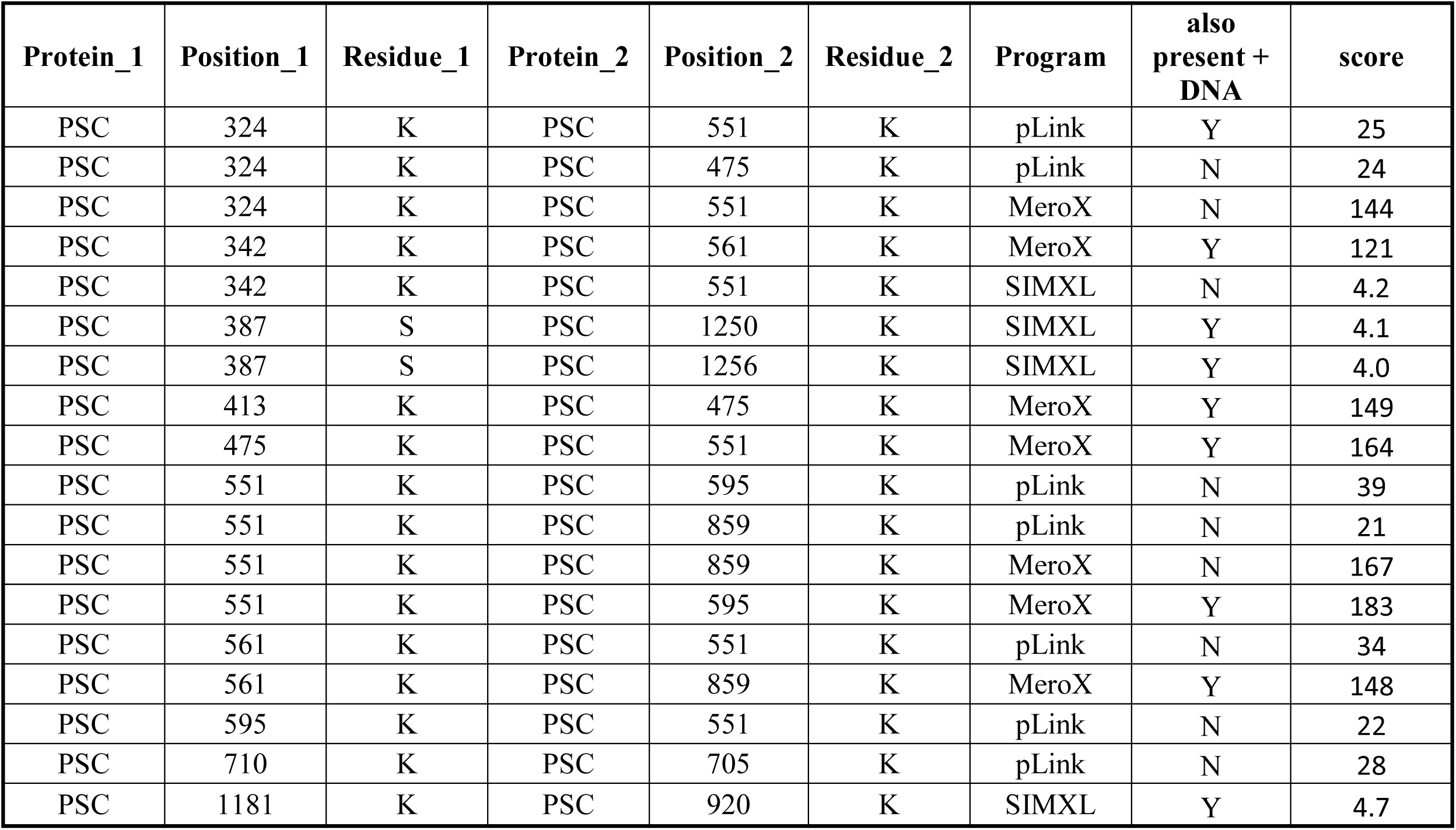
Intra-protein crosslinking sites in PSC in the absence of DNA. For each crosslink and each program, only the highest score obtained is stated. Score cutoffs are: 20 (pLink), 120 (MeroX), 4.0 (SIM-XL).

| **Protein_1** | **Position_1** | **Residue_1** | **Protein_2** | **Position_2** | **Residue_2** | **Program** | **also**<br>**present +**<br>**DNA** | **score** |
| --- | --- | --- | --- | --- | --- | --- | --- | --- |
| PSC | 324 | K | PSC | 551 | K | pLink | Y | 25 |
| PSC | 324 | K | PSC | 475 | K | pLink | N | 24 |
| PSC | 324 | K | PSC | 551 | K | MeroX | N | 144 |
| PSC | 342 | K | PSC | 561 | K | MeroX | Y | 121 |
| PSC | 342 | K | PSC | 551 | K | SIMXL | N | 4.2 |
| PSC | 387 | S | PSC | 1250 | K | SIMXL | Y | 4.1 |
| PSC | 387 | S | PSC | 1256 | K | SIMXL | Y | 4.0 |
| PSC | 413 | K | PSC | 475 | K | MeroX | Y | 149 |
| PSC | 475 | K | PSC | 551 | K | MeroX | Y | 164 |
| PSC | 551 | K | PSC | 595 | K | pLink | N | 39 |
| PSC | 551 | K | PSC | 859 | K | pLink | N | 21 |
| PSC | 551 | K | PSC | 859 | K | MeroX | N | 167 |
| PSC | 551 | K | PSC | 595 | K | MeroX | Y | 183 |
| PSC | 561 | K | PSC | 551 | K | pLink | N | 34 |
| PSC | 561 | K | PSC | 859 | K | MeroX | Y | 148 |
| PSC | 595 | K | PSC | 551 | K | pLink | N | 22 |
| PSC | 710 | K | PSC | 705 | K | pLink | N | 28 |
| PSC | 1181 | K | PSC | 920 | K | SIMXL | Y | 4.7 |

**Supplementary Table 2.** Intra-protein crosslinking sites in PSC in the presence of DNA. For each crosslink and each program, only the highest score obtained is stated. Score cutoffs are: 20 (pLink), 120 (MeroX), 4.0 (SIM-XL).

| **Protein_1** | **Position_1** | **Residue_1** | **Protein_2** | **Position_2** | **Residue_2** | **Program** | **also**<br>**present -**<br>**DNA** | **score** |
| --- | --- | --- | --- | --- | --- | --- | --- | --- |
| PSC | 114 | K | PSC | 551 | K | MeroX | N | 127 |
| PSC | 324 | K | PSC | 551 | K | MeroX | Y | 133 |
| PSC | 342 | K | PSC | 561 | K | MeroX | Y | 130 |
| PSC | 342 | K | PSC | 475 | K | SIMXL | N | 4.3 |
| PSC | 387 | K | PSC | 1256 | K | SIMXL | Y | 4.3 |
| PSC | 387 | K | PSC | 1250 | K | SIMXL | Y | 4.3 |
| PSC | 413 | K | PSC | 475 | K | MeroX | Y | 146 |
| PSC | 475 | K | PSC | 551 | K | MeroX | Y | 168 |
| PSC | 475 | K | PSC | 595 | K | MeroX | N | 123 |
| PSC | 535 | K | PSC | 551 | K | MeroX | N | 130 |
| PSC | 535 | K | PSC | 475 | K | SIMXL | N | 4.8 |
| PSC | 551 | K | PSC | 595 | K | MeroX | Y | 189 |
| PSC | 551 | K | PSC | 574 | S | MeroX | N | 137 |
| PSC | 551 | K | PSC | 705 | K | MeroX | N | 136 |
| PSC | 551 | K | PSC | 619 | K | MeroX | N | 130 |
| PSC | 551 | K | PSC | 826 | K | MeroX | N | 120 |
| PSC | 561 | K | PSC | 859 | K | MeroX | Y | 129 |
| PSC | 595 | K | PSC | 867 | K | MeroX | N | 128 |
| PSC | 826 | K | PSC | 867 | K | MeroX | N | 121 |
| PSC | 859 | K | PSC | 886 | K | MeroX | N | 175 |
| PSC | 867 | K | PSC | 974 | K | MeroX | N | 130 |
| PSC | 867 | K | PSC | 859 | K | SIMXL | N | 4.1 |
| PSC | 1162 | K | PSC | 1166 | K | SIMXL | N | 4.6 |
| PSC | 1181 | K | PSC | 920 | K | SIMXL | Y | 4.6 |
| PSC | 1181 | K | PSC | 919 | S | SIMXL | N | 4.6 |
| PSC | 1181 | K | PSC | 830 | S | SIMXL | N | 4.5 |
| PSC | 1181 | K | PSC | 968 | S | SIMXL | N | 4.0 |
| PSC | 1239 | K | PSC | 1250 | K | pLink | N | 24 |
| PSC | 1504 | Y | PSC | 1556 | T | MeroX | N | 140 |
| PSC | 1565 | K | PSC | 859 | K | SIMXL | N | 4.2 |
| PSC | 1565 | K | PSC | 1438 | K | SIMXL | N | 4.2 |
| PSC | 1565 | K | PSC | 1554 | K | SIMXL | N | 4.1 |
| PSC | 1568 | S | PSC | 475 | K | SIMXL | N | 4.2 |
| PSC | 1568 | S | PSC | 1554 | K | SIMXL | N | 4.1 |
| PSC | 1569 | S | PSC | 475 | K | SIMXL | N | 4.2 |

**Supplementary Table 3.** Inter-protein crosslinking sites between dRING and PSC in the absence of DNA. For each crosslink and each program, only the highest score obtained is stated. Score cutoffs are: 20 (pLink), 120 (MeroX), 4.0 (SIM-XL). Distances are provided for crosslinks that fall within published structures (PDB 4R8P).

| Protein1 | Position_1 | Residue_1 | Protein2 | Position_2 | Residue_2 | Program | also present<br>+ DNA | score | distance,<br>Å |
| --- | --- | --- | --- | --- | --- | --- | --- | --- | --- |
| dRING | 22 | K | PSC | 1 | M | SIMXL | Y | 4.4 |  |
| dRING | 22 | K | PSC | 6 | S | SIMXL | N | 4.1 |  |
| dRING | 22 | K | PSC | 342 | K | pLink | Y | 33 | 17 |
| dRING | 22 | K | PSC | 342 | K | SIMXL | Y | 5.8 | 17 |
| dRING | 22 | K | PSC | 919 | K | SIMXL | Y | 5.7 |  |
| dRING | 77 | S | PSC | 551 | K | MeroX | N | 144 |  |
| dRING | 222 | S | PSC | 455 | K | SIMXL | Y | 5.5 |  |
| dRING | 222 | S | PSC | 1401 | K | SIMXL | N | 5.4 |  |
| dRING | 222 | S | PSC | 1509 | K | SIMXL | N | 4.1 |  |
| dRING | 225 | S | PSC | 455 | K | SIMXL | N | 5.4 |  |
| dRING | 225 | S | PSC | 1401 | K | SIMXL | N | 5.7 |  |
| dRING | 225 | S | PSC | 1509 | K | SIMXL | N | 4.1 |  |
| dRING | 228 | S | PSC | 455 | K | SIMXL | N | 4.3 |  |

**Supplementary Table 4.** Inter-protein crosslinking sites between dRING and PSC in the presence of DNA. For each crosslink and each program, only the highest score obtained is stated. Score cutoffs are: 20 (pLink), 120 (MeroX), 4.0 (SIM-XL). Distances are provided for crosslinks that fall within published structures (PDB 4R8P).

| **Protein1** | **Position_1** | **Residue_1** | **Protein2** | **Position_2** | **Residue_2** | **Program** | **also present - DNA** | **score** | **distance, Å** |
| --- | --- | --- | --- | --- | --- | --- | --- | --- | --- |
| dRING | 22 | K | PSC | 1 | M | SIMXL | Y | 4.7 |  |
| dRING | 22 | K | PSC | 7 | K | SIMXL | Y | 4.1 |  |
| dRING | 22 | K | PSC | 342 | K | SIMXL | N | 5.5 | 17 |
| dRING | 22 | K | PSC | 341 | Y | MeroX | N | 131 | 15 |
| dRING | 22 | K | PSC | 342 | K | pLink | Y | 20 |  |
| dRING | 22 | K | PSC | 919 | S | SIMXL | Y | 4.3 | 17 |
| dRING | 157 | K | PSC | 475 | K | SIMXL | N | 4.6 |  |
| dRING | 222 | S | PSC | 455 | K | SIMXL | Y | 4.8 |  |
| dRING | 225 | S | PSC | 455 | K | SIMXL | N | 4.5 |  |
| dRING | 332 | K | PSC | 551 | K | pLink | N | 25 |  |

**Supplementary Table 5.** Intra-protein crosslinking sites within dRING in the absence of DNA. For each crosslink and each program, only the highest score obtained is stated. Score cutoffs are: 20 (pLink), 120 (MeroX), 4.0 (SIM-XL).

| **Protein1** | **Position_1** | **Residue_1** | **Protein2** | **Position_2** | **Residue_2** | **Program** | **score** | **also**<br>**present +**<br>**DNA** |
| --- | --- | --- | --- | --- | --- | --- | --- | --- |
| dRING | 332 | K | dRING | 349 | K | pLink | 29 | Y |
| dRING | 357 | K | dRING | 349 | K | pLink | 31 | N |
| dRING | 418 | K | dRING | 421 | K | pLink | 23 | N |

**Supplementary Table 6.** Intra-protein crosslinking sites within dRING in the presence of DNA. For each crosslink and each program, only the highest score obtained is stated. Score cutoffs are: 20 (pLink), 120 (MeroX), 4.0 (SIM-XL). Distances are provided for crosslinks that fall within published structures (PDB 4R8P).

| **Protein1** | **Position_1** | **Residue_1** | **Protein2** | **Position_2** | **Residue_2** | **Program** | **also present - DNA** | **score** | **distance, Å** |
| --- | --- | --- | --- | --- | --- | --- | --- | --- | --- |
| dRING | 22 | K | dRING | 106 | S | MeroX | N | 140 | 30 |
| dRING | 22 | K | dRING | 106 | S | SIMXL | N | 4.3 | 30 |
| dRING | 22 | K | dRING | 107 | K | SIMXL | N | 4.4 | 29 |
| dRING | 77 | S | dRING | 88 | K | MeroX | N | 121 | 14 |
| dRING | 77 | S | dRING | 106 | S | MeroX | N | 131 | 30 |
| dRING | 80 | K | dRING | 91 | S | MeroX | N | 155 | 7 |
| dRING | 80 | K | dRING | 92 | K | MeroX | N | 189 | 10 |
| dRING | 332 | K | dRING | 349 | K | pLink | Y | 20 |  |
| dRING | 349 | K | dRING | 357 | K | MeroX | N | 153 |  |
| dRING | 357 | K | dRING | 349 | K | SIMXL | N | 4.1 |  |

## Footnotes

1 Abbreviations: Polycomb Group (PcG); Cross-linking mass spectrometry (XL-MS), Posterior Sex Combs (PSC); Posterior Sex Combs C-terminal region (PSC-CTR); Polycomb (Pc); Polycomb Repressive Complex 1 (PRC1); Polyhomeotic (Ph); Homology region (HR)

